# A Deep Learning Pipeline for Nucleus Segmentation

**DOI:** 10.1101/2020.04.14.041020

**Authors:** George Zaki, Prabhakar R. Gudla, Kyunghun Lee, Justin Kim, Laurent Ozbun, Sigal Shachar, Manasi Gadkari, Jing Sun, Iain D.C. Fraser, Luis M. Franco, Tom Misteli, Gianluca Pegoraro

## Abstract

Deep learning is rapidly becoming the technique of choice for automated segmentation of nuclei in biological image analysis workflows. In order to evaluate the feasibility of training nuclear segmentation models on small, custom annotated image datasets that have been augmented, we have designed a computational pipeline to systematically compare different nuclear segmentation model architectures and model training strategies. Using this approach, we demonstrate that transfer learning and tuning of training parameters, such as the composition, size and pre-processing of the training image dataset, can lead to robust nuclear segmentation models, which match, and often exceed, the performance of existing, off-the-shelf deep learning models pre-trained on large image datasets. We envision a practical scenario where deep learning nuclear segmentation models trained in this way can be shared across a laboratory, facility, or institution, and continuously improved by training them on progressively larger and varied image datasets. Our work provides computational tools and a practical framework for deep learning-based biological image segmentation using small annotated image datasets.

## Introduction

High-Content Imaging (HCI) uses automated liquid handling, image acquisition and image analysis to screen the biological effect of hundreds of thousands of perturbing agents, such as RNAi, CRISPR/Cas9, and chemical compounds, by measuring cellular phenotypic changes in microscopy-based assays of interest (1,2). Because most cell types possess only one centrally positioned nucleus, and the nucleus is used to identify individual cells, the precise and automated nuclear segmentation of nuclei stained with specific fluorescent dyes is the first essential analysis step in many HCI workflows (3). In addition, nuclei tend to have a regular shape, and can be easily visualized with fluorescent DNA stains, which are cheap and provide high contrast. Nevertheless, notable exceptions to these desirable properties exist: cell types and primary cells can have dramatically different nuclear sizes, there can be substantial differences in nuclear shape (such as lobulation in polymorphonuclear cells of the immune system), and cells can grow tightly packed and/or in colonies thus leading to adjacent or overlapping nuclei. This makes it challenging to develop, test and implement robust algorithms that can provide accurate and automated nuclear segmentation results for a wide range of cell types and in different experimental conditions, even using the same image acquisition platform.

In recent years, computational advances in deep Convolutional Neural-Networks (CNN) have produced models that match, and sometimes exceed, human levels of performance in a variety of classification, clustering, and segmentation tasks in biological image analysis (4–6). More specifically, several studies have described novel CNN-based approaches for the segmentation of nuclei from fluorescence microscopy images (7–13). In particular, some of these efforts focused on training a variety of CNN architectures with relatively large and publicly available fluorescent image datasets of cells and nuclei, with the goal of ensuring that trained models could be generalized to novel image datasets without the need to retrain them (11,12).

While the collection and distribution of pre-trained models and large datasets of annotated biological images is highly useful because it can in principle avoid the need for model training, less is known about the performance of pre-trained models on images of primary cells or cell types which possess nuclear shapes previously not “seen” by the pre-trained models, and which were acquired on different microscopes (9,14). In these cases, it would be useful to test what, if any, are the most effective steps for obtaining precise nuclear segmentation by training CNN models on relatively small sets of images, which are readily available in most laboratories (9,13,14). Finally, from an end-user standpoint, few guidelines exist on how pre-existing nuclear segmentation models can be trained or re-trained, fine-tuned, adapted and improved on new image datasets acquired on the same instrument (13).

To address these questions, we designed and implemented an end-to-end computational pipeline to quickly train and evaluate the performance of machine-learning-based nuclear segmentation algorithms. This pipeline was first used to generate preliminary nuclear labels for about 4,000 nuclei from 4 different cell types and a total of 10 images, which were then manually corrected in an interactive fashion, and used to train and evaluate the performance of different CNN-based architectures for nuclear segmentation. The pipeline was used to explore different training strategies for two pre-existing CNN-based image segmentation model architectures: Feature Pyramid Network-2-watershed (FPN2-WS) (15), and Mask R-CNN (MRCNN) (16). To mimic practical scenarios in the laboratory, we trained and applied these two models to segment nuclei in four cell types that have very different nuclear morphology and that were acquired at different magnifications on the same high-throughput imaging microscope. Our results describe a practical path for researchers to train different CNN architectures in a semi-automated manner and indicate that these deep learning models can be efficiently trained with few images to segment heteromorphic nuclei from microscopy images.

## Materials and Methods

### Cells

HCT116 and U2OS cells were obtained from ATCC (Manassas, VA, USA). MCF10A cells were a kind gift of Daniel Haber (Harvard University, MA, USA). Cells were grown in 384-well plates (CellCarrier Ultra 384, PerkinElmer, Waltham, MA), fixed in 4% PFA and stained with DAPI. Primary human eosinophils were obtained from healthy donors enrolled under NIH protocol NCT000090662 and purified as previously described (17). The protocol was approved by the Institutional Review Board of the National Institute of Allergy and Infectious Diseases (NIAID) at the National Institutes of Health (NIH). Eosinophils were plated in Poly-D-Lysine coated 384-well plates (CellCarrier Ultra 384, PerkinElmer, Waltham, MA), then fixed in 4% PFA and stained with DAPI. Cells were plated and stained with DAPI either in biological replicate wells (i.e., cells were plated on different plates on different days) for MCF10A and HCT116 cells, or technical replicate wells (i.e., cells were plated on the same plate, on the same day, but in different wells) for U2OS and primary eosinophils.

### Image Generation

Fluorescence microscopy images of 384-well plates were acquired on a CV7000S spinning disk confocal microscope (Yokogawa, Japan). For all cells, we used a 405 nm solid-state excitation laser, a 405/488/568/640 nm quad excitation dichroic mirror, a 568 nm emission dichroic mirror, a 445/45 nm bandpass emission mirror, and an sCMOS camera (2550 × 2160 pixels). For MCF10A and eosinophils, we used a 60X water objective (NA 1.2), z-sectioning every 0.5 microns, and camera binning set to 2X2 pixels. For U2OS cells we used a single z-plane acquisition with a 20X air objective (NA 0.75) and camera binning set to 1X1 pixel. For HCT116 cells we used a single z-plane acquisition with a 20X air objective (NA 0.75) and camera binning set to 2X2 pixels. Proprietary geometric and shading correction software was run on the fly during acquisition for all cell types. In the case of 3D image acquisition, images were maximally projected on the fly.

### Computational Pipeline

The computational pipeline was based on the Snakemake language (18), and developed as a reproducible workflow capable of evaluating any number of different nuclear instance segmentation models. The analysis workflow was designed to run either on High-Performance Computing (HPC) clusters or on a local server equipped with NVIDIA™ Graphics Processing Units (GPUs). Snakemake is a generic workflow management software, where individual computational tasks are chained together to form a pipeline. Each task is configured by a set of input files and parameters, by a call to a bash script or a Python script that accepts the input files and processes them, and by a set of output files generated by the task.

### Ground Truth Labels Generation

The semantic segmentation labels of nuclei from fluorescence microscopy images used both in training and testing of the segmentation models were generated semi-automatically in two steps. First, preliminary labels were automatically generated using either classical image processing techniques, e.g., seeded watershed (19) or existing, publicly available DL-models for nuclear segmentation. In particular, since nuclei of MCF10A cells displayed high contrast and good separation, ground truth labels (instance segmentation masks) for these cells were generated using a simple traditional nuclear segmentation pipeline in KNIME (20,21), which included thresholding, gaps filling, connected components, and manual separation of a few adjacent nuclei. A set of Python scripts was used to convert these preliminary nuclear labels images into a format compatible with the interactive, web-based image annotation editing tool Supervisely (22). An expert cell biologist corrected nuclear segmentation mistakes using a combination of bitmask brushes and polygons to generate a set of high-quality Ground Truth (GT) nuclear labels. As an example, for one of the tests datasets (e.g MCF10A_Biological, Semi-Automated, Table S1), the semi-automated correction task took about 2 hours for 1,349 nuclei from 7 images. As a control, we also generated a test dataset from the same images in which the nuclei were fully manually annotated to assess for possible biases introduced by the semi-automated ground-truth generation strategy. The manual annotation was carried on using bitmasks brushes in Supervisely. As an example of one of these test datasets (e.g. MCF10A_Biological, Manual, Table S1), the fully manual labelling of nuclei from these images took approximately 12 hours for annotating 1,382 nuclei.

The nuclear GT labels for MCF10A cells were used for a first round of supervised training for the two deep learning models used in this study, MRCNN and FPN2-WS (see below for details). The DL models trained in this fashion were then used in inference mode to generate preliminary labels for all other cell types, which were then similarly corrected using Supervisely to obtain the semi-automated GT labels. At every step of this process, the manual annotator was blinded with respect to the algorithm/model that produced the preliminary nuclear labels.

### Training/Validation/Testing Strategy

Images and GT labels for the cell types defined above were divided into two sets: training and testing (Table S1). For every given cell type, the training datasets (MCF10A_Original, U2OS_Original, HCT116_Original, and Eosinophils_Original) (Table S1) were used in both model training and model validation. When we trained the models on training datasets from all cell types, as reported in Fig. 5, we had a total of 4,012 nuclei in 10 full Fields of View (FOVs) (Table S1). After random sampling of Regions of Interest (ROIs) from these FOV and ROI augmentation (for a total of 35,000 ROI’s for most experiments, see Image Augmentation subsection below), we assigned 80% of the total number of augmented ROIs for model training and assigned the remaining 20% for model validation (see Model Training subsection). The testing datasets (MCF10A_Biological, U2OS_Technical, HCT116_Biological, and Eosinophils_Technical) (Table S1) were used only for assessing the performance of the DL models at inference. We had a total of 3,607 nuclei in 10 FOVs for the testing datasets from all cell types (Table S1). All the F1 scores reported in Fig. 2–5, Tables 1 – 8, and Fig. S2A were calculated using the testing datasets and semi-automated GT labels. F1 scores in Fig. S1B and Table S3 were calculated using testing datasets and both manual and semi-automated GT labels. F1 scores in Fig. S2B and Table S4 were calculated using both training and testing datasets and semi-automated GT labels.

**Table 1).** F1 Scores at IoU[t = 0.7] and IoU[t = 0.9] for the baseline strategy for the MRCNN and FPN2-WS models. MRCNN and FPN2-WS were trained only on the MCF10A cell line training image dataset using random initialization, as indicated in Fig. 2. The table indicates the test F1 scores for MRCNN and FPN2-WS object-level segmentation (IoU[t = 0.7]) and pixel-level segmentation (IoU[t = 0.9]) for each 4 cell types. The best F1 value for each IoU/Cell Type combination is indicated in bold.

| Model | MCF10A/o.7 | MCF10A/o.9 | HCT116/o.7 | HCT116/o.9 | U2OS/o.7 | U2OS/o.9 | Eosinophils/o.7 | Eosinophils/o.9 |
| --- | --- | --- | --- | --- | --- | --- | --- | --- |
| FPN2-WS | 0.97 | 0.97 | 0.84 | 0.2 | 0.94 | 0.92 | 0.48 | 0.21 |
| MRCNN | 0.82 | 0.75 | 0.63 | 0.07 | 0.84 | 0.65 | 0.72 | 0.06 |

### Image Augmentation

Image augmentation of FOVs was applied to training datasets to simulate slightly different illumination conditions, to enrich the training image dataset, and to avoid model overfitting. Multiple augmentation methods from the Python module *imgaug* (23) were used. In particular, our pipeline can be configured to use scaling transformations (0.33X, 0.66X, 1X, 1.33X, and 2X), affine transformations (vertical and horizontal flips, 45°, 90°, 135°, 180°, 225°, 270°, and 315° rotations), contrast transformations (Contrast Limited Adaptive Histogram Equalization (CLAHE) (24), gamma correction, and saturation), blurring (median blur and gaussian blur), or random noise addition (impulse noise, gaussian noise, and dropout).

When only MCF10A cells were used for model training, we first applied the 5 scaling transformations to the FOVs and then randomly sampled 7,000 overlapping ROIs of size 256 × 256 pixels from each set of scaled FOVs of the grayscale images and from the matching ground truth labels images, resulting in a total of 35,000 augmented ROIs. In cases when all four cell types were used for model training, each cell type contributed 8,750 scaled and augmented ROIs, again for a total of 35,000 ROIs (With the exception of Fig 4B, where different numbers of ROIs were used for model training). For each scaled ROI, one of the affine transformations was randomly picked and applied. For each scaled and rotated or flipped ROI, one of the intensity augmentation methods mentioned above was randomly selected with a probability between 0 and 1 and applied to the grayscale ROI and to the nuclear label ROI, when appropriate. For the MRCNN architecture, the augmented ground truth labels were converted to binary mask images. For the FPN2-WS architecture, the labels were further transformed using traditional image processing operations (contour generation, distance transform) to obtain the normalized distance transform and gaussian blurred grayscale images used in this model (See below for further details). Augmented ROIs from all the cell types were combined and saved in HDF5 (.h5) file format.

In experiments where the image augmentations were turned off, we assigned a probability of 0 to the augmentation classes to disable them. Rotations and flips were applied in all training experiments.

### Model Description

#### Mask R-CNN (MRCNN)

We used the Matterport implementation of Mask R-CNN (MRCNN) (25) with minor modifications. In particular, we added a parser to accept images in .h5 format and set the size of the input layer for both model training and inference to 256 × 256 pixels. At inference time, full FOVs were first divided into overlapping grids of 256 × 256 pixels, where the overlap value was set to 50 pixels, which on average was enough to cover one nucleus. Model inference was then run on the ROIs for a single FOV to generate nuclear labels with a unique ID. Nuclear segmentation labels with a different ID that were overlapping above a certain threshold across two or more ROIs were then merged into a nuclear label with a single ID. Nuclear labels with an area below a certain threshold were eliminated. Finally, since MRCNN can sometimes predict more than one connected component per nuclear label ID, only the largest connected component for every nuclear label ID was retained.

#### Feature Pyramids Network-2-watershed (FPN2-WS)

In the FPN2-WS architecture, two feature pyramids networks (FPN) (26) based models, each one with one output layer, were trained to predict a normalized distance transform image for each nucleus and a gaussian blurred (σ = 1) version of the nuclear label border, respectively. In the normalized distance transform the pixels values ranged between 0 and 1, such that pixels closer to the nuclear boundary have values closer to 0 and pixels at the center of the nucleus are closer to 1. The predicted normalized distance transform the FPN2-WS model was then thresholded at a value of 0.6. The thresholded normalized distance transform and the blurred border images were used as inputs to the seeded watershed segmentation algorithm (19) for delineating nuclei.

### Model Training

For all 4 cell types, full FOV of DAPI-stained cells from a single replicate well were used as a training dataset (Table S1). One training cycle was defined as a single pass of the entire augmented dataset through the network. We decided to use the term “cycle”, as opposed to the more traditional term “epoch”, because the original images were randomly bootstrapped to generate a training dataset of 35,000 256 × 256 pixels ROIs. Unless otherwise stated, the MRCNN and FPN2-WS models were trained for a total of 25 cycles. The model with the lowest validation loss value over 25 cycles was saved and used for inference. The model parameters are available in a dedicated Figshare project repository (28).

For MRCNN, we used the same learning rate schedule as previously described in the MatterPort implementation of this model (25). In particular, since the MatterPort schedule is defined over 200 iterations, we then distributed the 25 training cycles over 200 iterations starting with a learning rate of 10e-4.

For the FPN2-WS model, in training experiments involving initializing the network with pre-trained weights, e.g., ImageNet, the network parameters/weights of the encoder portion were frozen during training for the first 2 cycles. After the two warmup cycles, all trainable layers were unfrozen and continued training for the remaining 23 cycles. We selected a sigmoid activation function, and the Adam optimizer to minimize the mean square error of the corresponding network output image (i.e., distance transform or blurred contour, respectively). The learning rate was set to a constant value of 1e-3. All the training experiments are described in Table S2, where for each experiment the initialization, training datasets, number of training cycles, number of ROIs, and augmentation strategies are indicated.

### Model Testing

For inference, each model was tested on images of nuclei from all cell types obtained either from technical (U2OS, eosinophils) or biological replicate wells (MCF10A, HCT116) that were not used for model training, and, unless otherwise stated, semiautomated GT labels were used for scoring (See Table S1). In Fig. S2B we tested model performance on both training and testing image datasets with their associated semiautomated GT labels to estimate potential model overfitting.

### Quantitative Assessment of Nuclear Instance Segmentation Performance

Given ground truth nuclei IDs and the inference nuclei IDs, the F1 scores were calculated at different Intersection over Union (IoU) thresholds. A two-dimensional histogram was used to calculate the intersection area (i.e., the number of pixels) between all pairs of nuclei from the ground truth labels and the inference labels. The union area between every pair of ground truth and inference labels was calculated as the sum of the pixels belonging to either of these labels minus their intersection. For every pair of ground truth and inference labels, the IoU was calculated as the ratio between the intersection over the union. For any IoU threshold *t* between 0 and 1, the True Positive (*TP*(*t*)) statistic was the number of ground truth labels that have an IoU value equal or higher than *t* with one inference label. Similarly, the False Negative (*FN*(*t*)) statistic was the number of ground truth labels that have an IoU value lower than *t* with all inference labels. Finally, the False Positive (*FP*(*t*)) was the number of inference labels that have an IoU value lower than *t* with all ground truth label.

Given *TP*(*t*), *FN*(*t*) and *FP*(*t*), the F1 score at a given IoU threshold *t* was calculated as:

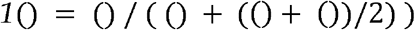

Furthermore, an over-splitting event was defined as a type of error where a nucleus in GT is matched with more than one inference nucleus with an IoU of at least 0.1. A merge error was defined as a type of error where an inference label was matched to more than one GT label with an IoU of at least 0.1.

The IoU used for calculating F1 score can aggressively penalize the performance readout of segmentation models for datasets with predominantly smaller objects (e.g., circular objects with a radius of 5 pixels). The penalty is more severe at threshold values *t* > 0.6. To illustrate this, consider one dataset in which all objects are perfect circles with radius 60 pixels and another dataset in which all objects are also perfect circles with radius of 4 pixels. These two datasets, with some approximations, simulate images acquired on an optical microscope using 4X and 60X objectives. Also assume that the segmentation model always makes one-pixel error along the boundary on both datasets. It can be seen that for the dataset with objects with a radius of 60 pixels the IoU will be 0.967 whereas for the datasets with objects with a radius of 4 pixels the IoU will be 0.563. Thus, at an IoU threshold value t=0.6 (IoU[t = 0.6]) the segmentation model will appear to be severely underperforming.

### Software Repository

The Snakemake pipeline for models training and testing, the weights for the best MRCNN and FPN2-WS models, the images used, and the code used to generate Figs. 2 – 5, Figs. S1 – S3, Tables 1 – 8, and Tables S3 and S4 are deposited in a dedicated Github repository (27), and in a dedicated Figshare project repository (28).

## Results

### A Practical Framework for Training and Testing Deep Learning Nuclear Segmentation Models on Small Annotated Image Datasets

We sought to test the practical implementation of different Convolutional Neural Networks (CNN) image segmentation models and training strategies for the automated identification of nuclei from a wide variety of fluorescence microscopy images. To this end, we built a computational pipeline (See Materials and Methods for details) to train the CNN models starting from a limited number of fluorescence microscopy images (1 to 7 Fields of View (FOVs) per cell line, depending on the cell line; Table S1), which is a typical use case scenario for a single cell biologist/researcher. As a first step in the pipeline, a pre-trained CNN model or traditional nucleus segmentation approaches were used to generate preliminary sets of labels for the nuclei (Fig. 1A). Depending on a variety of factors, these preliminary labels contained gross segmentation mistakes, such as false positives, false negatives, splits and merges, which had to be manually corrected using an interactive, web-based interface (Fig. 1A, and Materials and Methods).

**Fig. 1).**
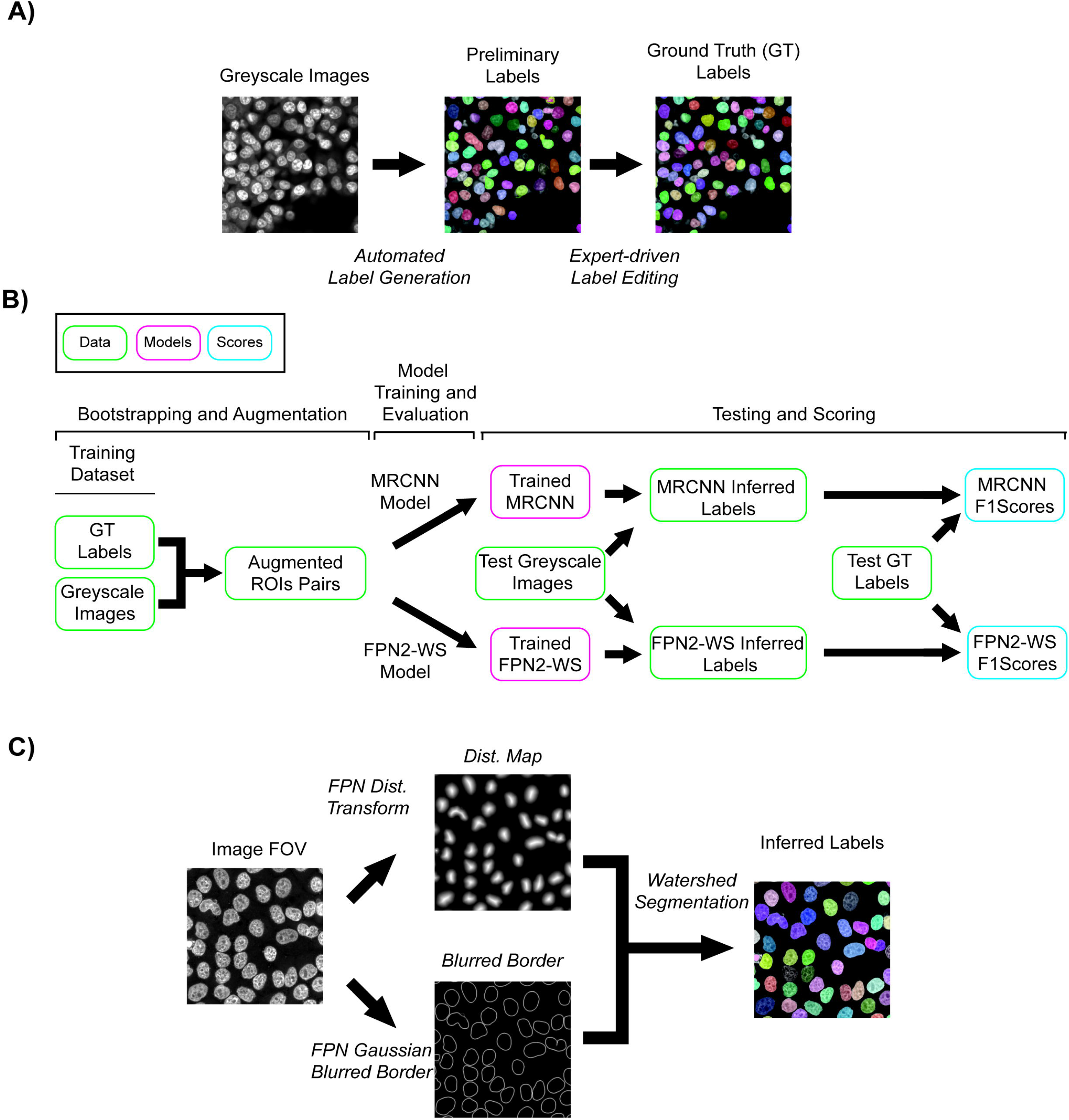
Deep Learning nuclear segmentation pipelines. **A)** Schematic representation of the semi-automated approach to generate Ground Truth (GT) labels. Either a traditional image processing algorithm, or a pre-trained CNN segmentation model were used to generate a set of preliminary labels for the nuclei. These were manually corrected using a web-based interface to generate a set of GT nuclear labels. **B)** Workflow of the CNN model training and testing strategies. Random overlapping regions of interest (ROIs) are oversampled (bootstrapped) from full fields of view (FOVs) of grayscale images and of GT labels images. These ROIs were then augmented by randomly applying a variety of affine, translational, intensity and blurring transformations. The augmented dataset of ROIs is then split 80%-20% for training and validation, respectively, of the deep learning nuclear segmentation models. Trained models are then tested by running inference on a set of images not seen at training or validation. Test model performance is scored using the corresponding GT labels for the test dataset and F1 scores are generated. **C)** Schematic of the Feature Pyramid Network-2-watershed (FPN2-WS) model architecture. FPN2-WS uses two deep learning models to predict a normalized nucleus distance transform image, and a blurred nucleus border image, respectively. These images are then used as the seed and border for a traditional seeded watershed segmentation algorithm.

This set of semi-automatically generated ground-truth (GT) labels was then used to train CNN models in combination with the original grayscale images of DAPI-stained nuclei and to test their performance (Fig. 1B). A set of images and their associated GT labels per each cell line was set aside for model training and validation (Fig. 1B and Table S1). Since we decided to train the deep learning segmentation models from relatively small datasets of images, we decided to bootstrap the datasets by generating multiple random overlapping regions of interest (ROIs) from the original full FOVs and to augment these ROIs by randomly applying several image transformations (Fig. 1B and Materials and Methods for details). Following these steps, the CNNs were trained with an equal number of augmented ROIs for each cell line (Fig. 1B) (23). In addition, in most model training regimens, and to speed up training and to ensure that the models would reach convergence during training, we also adopted a transfer learning approach (29,30), where the models were initialized with weights obtained from training on large datasets of everyday images, such as ImageNet (31) or COCO (32) (Fig. 1B). Finally, the nuclear segmentation performance of trained models was tested at inference by calculating F1 scores at different Intersection over Union (IoU) thresholds for a dedicated test set of greyscale images and GT labels that were never used for model training (11) (Fig. 1B, Table S1, and Materials and Methods for details).

To test the training pipeline for deep learning nuclear segmentation on different CNN architectures, we trained two models widely used in instance segmentation tasks: Feature Pyramid Network-2-watershed (FPN2-WS) (15), and Mask-RCNN (MRCNN) (16). In particular, and similarly to previously described approaches (11,33), FPN2-WS (Fig. 1C) consisted of two separate FPN models to predict a normalized distance transform image for the nucleus and a Gaussian blurred image of its border, respectively (Fig. 1C). These two images were then thresholded and used as the seed and border, respectively, in the seeded watershed segmentation algorithm (19) to obtain the final nuclear labels (Fig. 1C).

### Testing performance of randomly initialized models trained on images of nuclei from a single cell line

With this training/testing framework in place, we proceeded to test several different training strategies (Table S2). Our baseline for training consisted of randomly initialized MRCNN and FPN2-WS models, trained using 35,000 randomly cropped and augmented ROIs of 256 × 256 pixels from 7 FOVs containing 1,225 nuclei from the epithelial MCF10A cell line, which has a regular nuclear morphology and, for the most part, does not form clumps/colonies or occlusions (Fig. 2A, Table S1). We then tested the inference performance of the trained MRCNN and FPN2-WS models on previously unseen test datasets: (i) MCF10A nuclei from another biological replicate imaged at 60X magnification, (ii) osteosarcoma cell line U2OS nuclei from another technical replicate imaged at 20X, (iii) colorectal cell line HCT116 nuclei from another biological replicate imaged at 20X, and (iv) primary human eosinophils nuclei from another technical replicate imaged at 60X (Fig. 2A and Table S1). These cell types were chosen to test the performance of the trained nuclear segmentation architectures on a range of different nuclear shapes, sizes, confluencies, and occlusions (Fig. 2A). Not surprisingly, visual comparison of the GT nuclear labels (Fig. 2A, GT panels), with the nuclear labels predicted by the MRCNN and FPN2-WS trained as indicated above (Fig. 2A, Inference panels), revealed the presence of several prediction errors (Fig. 2A, GT XOR Inference panels). These included instances of under-segmentation, over-segmentation, missed nuclei, and erroneous nuclear borders. These errors were particularly evident in HCT116 cells, which tend to grow in clumps, and in primary eosinophils, which have a highly lobulated nucleus (Fig. 2A). Accordingly, quantification of these images by measuring F1 inference scores at an IoU threshold of 0.7 (IoU[t = 0.7], see Materials and Methods for details) (11), showed that MRCNN had reasonably high F1 score values for MCF10A, U2OS and lower ones for HCT116, and primary eosinophils (Fig. 2B and Table 1), while the FPN2-WS model had high F1 score values for MCF10A, U2OS, and HCT116, but fairly low scores for primary eosinophils, respectively (Fig. 2B and Table 1). The results of this first set of experiments suggest that training CNN models with random initialization using a single cell type does provide, at least in the case of FPN2-WS, good nuclear segmentation performance on the same cell type, or on cells with similar nuclear morphology and confluency characteristics (Fig. 2A, compare MCF10A with U2OS). At the same time, the models failed to generalize at inference time on other cell types with more heteromorphic nuclei or growth characteristics (Fig. 2A, compare MCF10A with HCT116 and primary eosinophils).

**Fig. 2).**
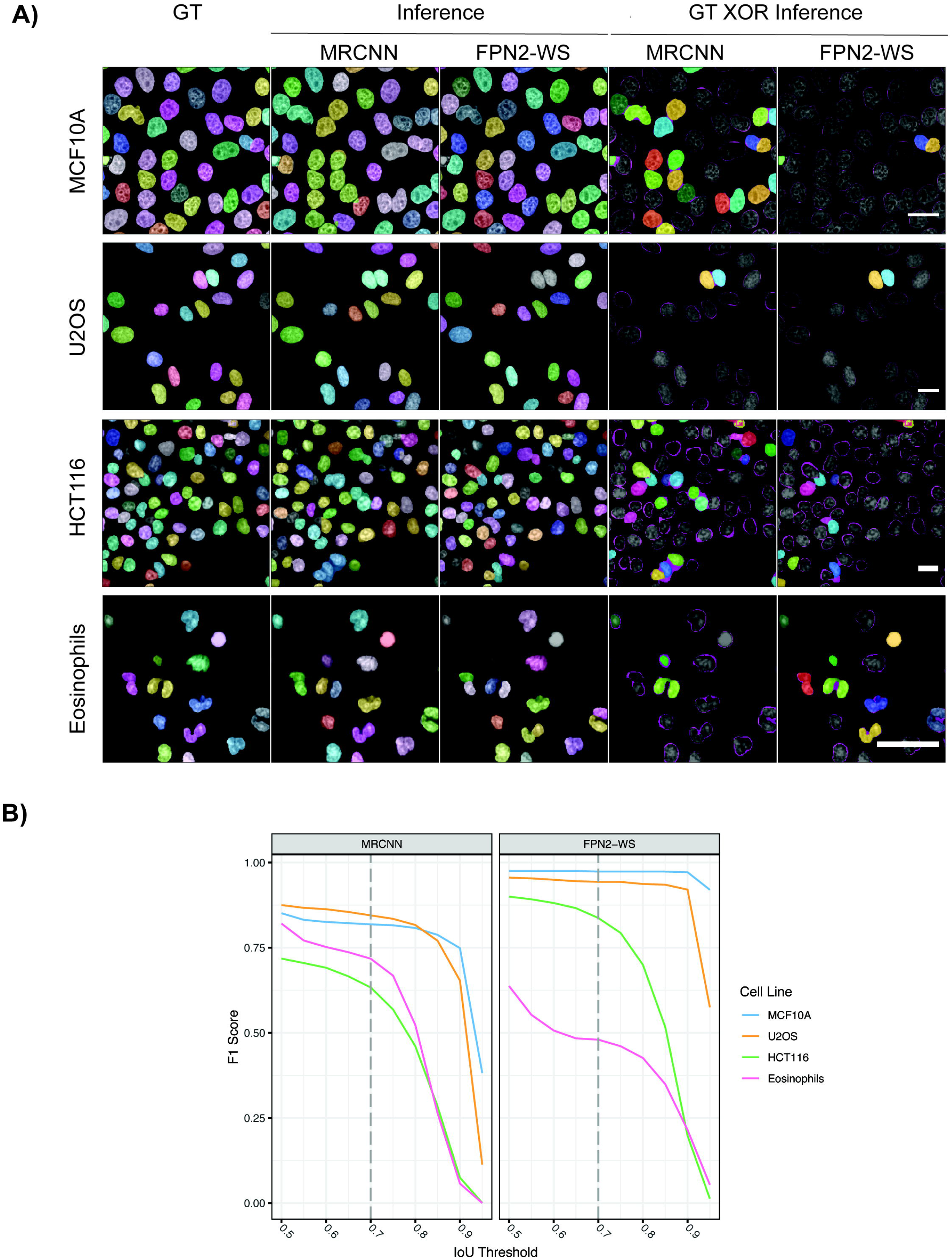
Nuclear segmentation inference performance of the baseline training strategy for the MRCNN and FPN2-WS model architectures. **A)** The MRCNN and FPN2-WS models were trained on the MCF10A cell line image dataset (35,000 bootstrapped and augmented ROIs) using random initialization, and applied inference on test images of nuclei from the indicated cell types. The images represent pseudocolored nuclear labels for the ground truth (GT) annotations and for the model inference results on test images. Additionally, the GT XOR Inference panels represent pseudocolored full labels for false negatives at an Intersection over Union (IoU) threshold of 0.7 (IoU[t = 0.7]). For true positive and false positive objects at the same threshold, only the difference at the pixel-level between GT and inference labels is shown in magenta. Scale bar: 20 μm. **B)** Line plot of the test F1 Score at increasing Intersection over Union (IoU) thresholds (See Materials and Methods for details) for the MRCNN and FPN2-WS models trained as indicated in A). IoU[t = 0.7] is indicative of object-level segmentation performance on test images and is indicated by the dashed grey vertical line in the plot.

### Improvement of Model Segmentation Performance by Transfer Learning and Progressive Enrichment of Training Datasets with Images of Nuclei with Diverse Morphology

We thus tested whether changing the model training strategy would improve the performance of these deep learning models on our image inference datasets. In particular, to avoid an excessive increase in the number of values and parameters to be tested, we decided to not change CNN network architectures, the training optimizing algorithms, or the learning rate schedule. Instead, we focused on varying the CNN weights initialization (i.e., transfer learning), the composition of the image training dataset, the number of training cycles, the size of the training dataset, and the augmentation strategy.

We began testing the effect of these changes by initializing the MRCNN and FPN2-WS models with weights obtained by training them on the COCO (32) or ImageNet (31) datasets, respectively (Fig. 3A and Table 2). As compared to random initialization, pre-initialization with COCO weights and transfer learning increased F1 inference scores at IoU[t = 0.7] of the MRCNN model on the MCF10A, U2OS, and HCT116 cell types, while leading to a decrease for the primary eosinophils (Fig. 3 A and Table 2). In similar experiments using the FPN2-WS model, transfer learning had a marginal increase of F1 scores on MCF10A, U2OS and HCT116, and a moderate increase of the F1 score in eosinophils (Fig. 3A and Table 2).

**Fig. 3).**
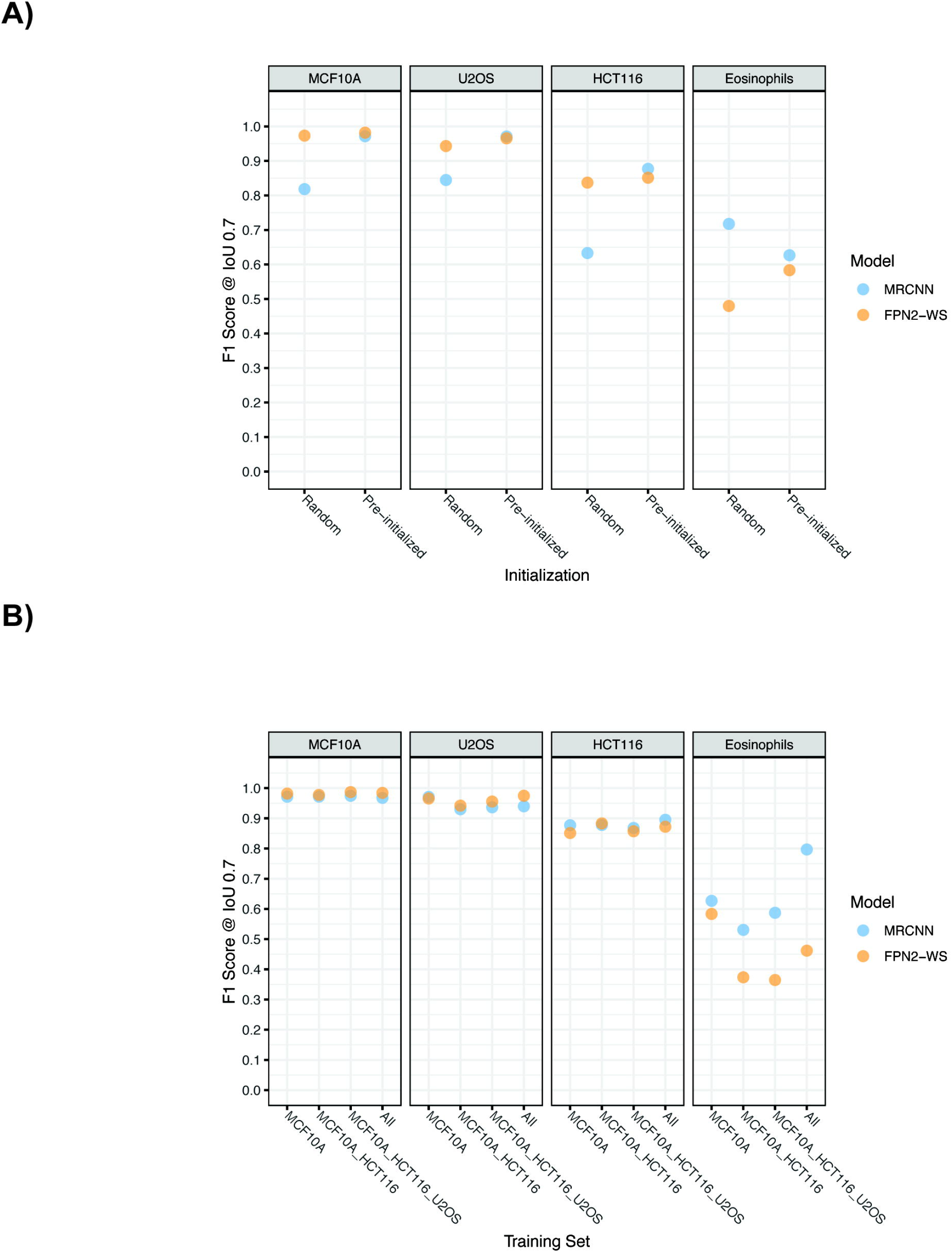
Transfer learning can improve the inference performance of nuclear segmentation models. **A)** Dot plot of the test F1 Score at IoU[t = 0.7] indicative of object-level segmentation performance for the nuclear segmentation models trained either as indicated in Fig. 1A) (35,000 bootstrapped and augmented ROIs obtained from the MCF10A cell line training dataset were used, Randomized weights), or by using the same training data and by initializing the models with weights obtained by pre-training MRCNN on the COCO dataset, or the FPN2-WS on the ImageNet dataset, respectively. **B)** Dot plot of the test F1 Score at IoU[t = 0.7] for the nuclear segmentation models trained either as indicated in Fig. 3A) (35,000 bootstrapped and augmented ROIs obtained from the MCF10A cell line training dataset were used, pre-initialized weights), or by using a combination of equal numbers of bootstrapped and augmented ROIs from the training datasets of other cell types, as indicated on the x-axis, for a total of 35,000 ROIs in each case.

**Table 2).** F1 Scores at IoU[t = 0.7] for MRCNN and FPN2-WS models with different pre-initialization strategies. MRCNN and FPN2-WS were trained only on the MCF10A cell line training image dataset using random initialization or transfer learning with pre-trained model weights, as indicated in Fig. 3A. The table indicates the test F1 scores for MRCNN and FPN2-WS object-level segmentation (IoU[t = 0.7]) for each 4 cell types in the training conditions examined. The best F1 value for each Cell Type/Model/Pre-initialization combination is indicated in bold.

| Model | Initialization | MCF10A | HCT116 | U2OS | Eosinophils |
| --- | --- | --- | --- | --- | --- |
| FPN2-WS | Random | 0.97 | 0.84 | 0.94 | 0.48 |
| FPN2-WS | Pre-initialized | **0.98** | 0.85 | 0.97 | 0.58 |
| MRCNN | Random | 0.82 | 0.63 | 0.84 | **0.72** |
| MRCNN | Pre-initialized | 0.97 | **0.88** | **0.97** | 0.63 |

Having observed a positive effect of transfer learning on nuclear segmentation tasks using these models, we then progressively increased the variety of images used in training of pre-initialized models to test whether these are capable of learning a set of nuclear segmentation features for multiple cell types in sequential or simultaneous fashion (Fig. 3B and Table 3). At the end of this process, 35,000 augmented ROIs from a total of 10 FOVs of the 4 different cell types for a total of 4,082 nuclei were used in training (Table S1). The F1 scores at IoU[t = 0.7] for MCF10A, U2OS and HCT116 nuclei did not appreciably increase or decrease upon training MRCNN and FPN2-WS with data from multiple cell types (Fig. 3B and Table 3). On the other hand, for the primary eosinophils, the F1 score for MRCNN increased when images from these cells were used in training, as compared to when only MCF10A cells were used (Fig. 3B and Table 3).

**Table 3).** F1 Scores at IoU[t = 0.7] for the MRCNN and FPN2-WS models trained on increasingly diverse image datasets. MRCNN and FPN2-WS were trained only on the MCF10A cell line training image dataset, or on progressively more complex image datasets, as indicated in Fig. 3B. The table indicates the test F1 scores for MRCNN and FPN2-WS object-level segmentation (IoU[t = 0.7]) for each 4 cell types in the training conditions examined. The best F1 value for each Cell Type/Model/Image Training Set combination is indicated in bold.

| Model | Training Set | MCF10A | HCT116 | U2OS | Eosinophils |
| --- | --- | --- | --- | --- | --- |
| FPN2-WS | MCF10A | 0.98 | 0.85 | 0.97 | 0.58 |
| FPN2-WS | MCF10A-HCT116 | 0.98 | 0.88 | 0.94 | 0.37 |
| FPN2-WS | MCF10A-HCT116-U2OS | **0.99** | 0.86 | 0.96 | 0.36 |
| FPN2-WS | All | 0.98 | 0.87 | **0.97** | 0.46 |
| MRCNN | MCF10A | 0.97 | 0.88 | 0.97 | 0.63 |
| MRCNN | MCF10A-HCT116 | 0.97 | 0.88 | 0.93 | 0.53 |
| MRCNN | MCF10A-HCT116-U2OS | 0.97 | 0.87 | 0.94 | 0.59 |
| MRCNN | All | 0.97 | **0.9** | 0.94 | **0.8** |

These results indicate that using pre-initialization of models with weights from large image datasets has a positive effect on nuclear segmentation using the MRCNN model on most types tested. In addition, sequential training of deep learning models with increasingly varied image training sets does not have a negative impact on the performance of the deep learning models tested here, and in the case of primary eosinophils, which are polymorphonuclear cells, it can improve the performance of MRCNN.

### Analysis of Training Parameters Important for Nuclear Segmentation Performance

Image pre-processing, such as aggressive image augmentation, can become a time and computing power bottleneck if the DL model training step is relatively fast. In addition, DL models can require longer training times if the training dataset is large. We thus considered the impact of reducing the number of training cycles (Fig. 4A and Table 4), or the number of augmented image ROIs in training datasets for pre-initialized MRCNN or FPN2-WS models trained on images from all the cell types (Fig. 4B and Table 5). Reducing the number of training cycles by a factor of 4 (Fig. 4A and Table 4, 1X vs 0.25X) led to a slight decrease in F1 inference scores at IoU[t = 0.7] for MCF10A, U2OS and HCT116, while it resulted in a larger decrease for primary eosinophils for both MRCNN and FPN2-WS models. Similarly, to test whether we could maintain the same F1 inference scores at IoU[t = 0.7] by training the models with fewer ROIs, or if we could improve them by training with more ROIs, we trained pre-initialized MRCNN and FPN2-WS with ROIs from images of all the 4 cell types, ranging from 8,750 to 70,000 to augmented ROIs, reflecting a 8-fold range of training dataset sizes (Fig. 4B and Table 5, 0.25X to 2X) as compared to the number of training ROIs used in the previous experiments (1X, 35,000 augmented image ROIs) (Fig. 4B and Table 5). Training with twice the original number of ROIs did not lead to an improvement in F1 inference scores for either model in any of the cell types (Fig. 4B and Table 5). Reducing the number of ROIs 4-fold from the original training dataset instead had more diverse effects on the F1 inference scores, ranging from negligible (e.g. Fig 4B and Table 5, MCF10A and MRCNN) to substantial decreases (e.g. Fig 4B and Table 5, eosinophils and MRCNN). Finally, we tested the effect of eliminating one class of augmentation operations at a time, or all at the same time, to test the contribution of each type of augmentation on the performance of the model (Fig. 4C and Table 6). Surprisingly, for both MRCNN and FPN2-WS, and as compared to implementing the full set of augmentations, skipping all image augmentations, with the exception of random rotations and flips, did not degrade the F1 inference performance at IoU[t = 0.7] of the models for any cell type (Fig. 4C and Table 6) and slightly improved F1 inference scores for MRCNN in the case of U2OS cells, HCT116 and primary eosinophils.

**Fig. 4).**
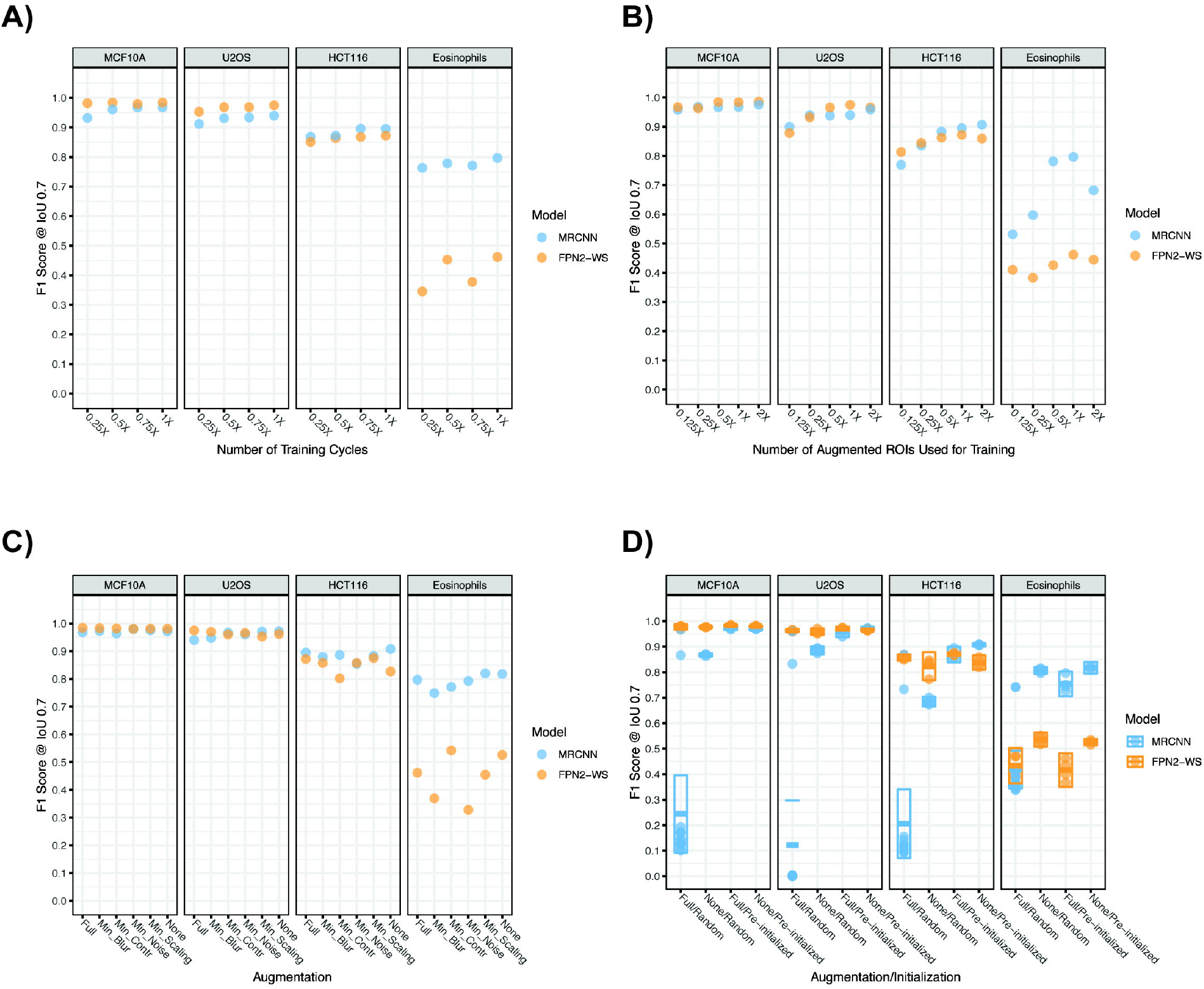
Deep learning segmentation models can be efficiently trained with less data, fewer training cycles, and fewer image pre-processing steps. **A)** Dot plot of the test F1 Score at IoU[t = 0.7] indicative of object segmentation performance for the nuclear segmentation models trained with images of nuclei from all cell types as indicated in Fig. 3B) (1X, 25 cycles, 35,000 bootstrapped and augmented ROIs obtained from all the cell types training datasets, pre-initialized weights), or by reducing the number of training cycles and keeping the other training parameters constant (0.25 - 0.75X, 6.25 - 18.75 cycles, See Materials and Methods for the definition of training cycles). **B)** Dot plot of the test F1 Score at IoU[t = 0.7] for the nuclear segmentation models trained either from all cell types as indicated in Fig. 3B) (1X, 35,000 bootstrapped and augmented ROIs obtained from all the cell types training datasets), twice as many ROIs (2X, 70,000 total), or fewer ROIs (0.125X - 0.5X, 4,375 - 17,500 total). **C)** Dot plot of the test F1 Score at IoU[t = 0.7] for the nuclear segmentation models trained on 35,000 bootstrapped ROIs obtained from all the cell types training datasets that were only augmented by ROI rotations and flips (None), or were augmented with all but one of the other selected class of augmentation operations (Min_Blur - Min_Scaling) as indicated on the x-axis. **D)** Dot plot and crossbar plot for the mean and 95% confidence interval crossbar for the test F1 Score at IoU[t = 0.7]. Nuclear segmentation models were independently trained with 35,000 bootstrapped ROIs obtained from all the cell types training datasets, for 25 cycles, and as indicated on the x-axis. n = 15 for the MRCNN Full/Random combination, and n = 4 for all other combinations.

**Table 4).**
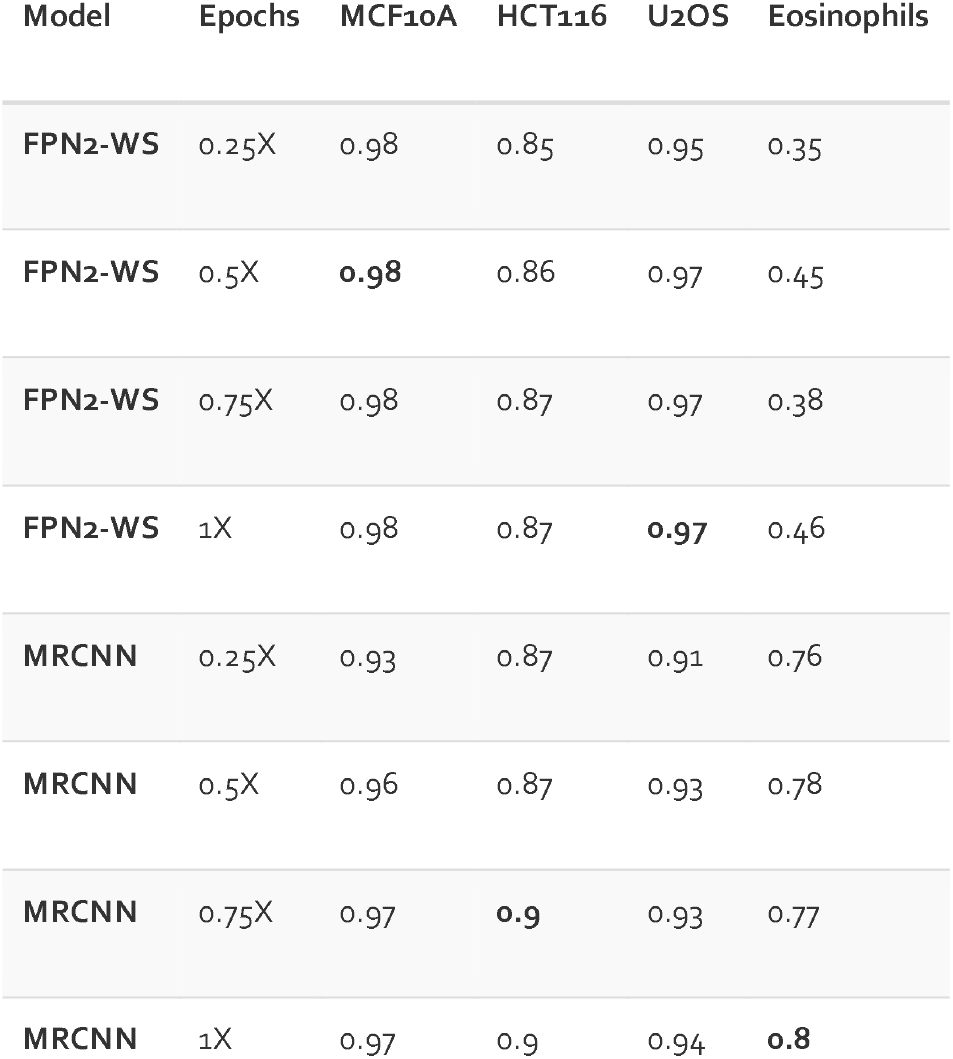
F1 Scores at IoU[t = 0.7] for the MRCNN and FPN2-WS models trained for an increasing amount of times. MRCNN and FPN2-WS were trained only on image datasets from all cell types training image dataset for increasing number of epochs, as indicated in Fig. 4A. The table indicates the test F1 scores for MRCNN and FPN2-WS object-level segmentation (IoU[t = 0.7]) for each of the 4 cell types in the training conditions examined. The best F1 value for each Cell Type/Model/Training length combination is indicated in bold.

| Model | Epochs | MCF10A | HCT116 | U2OS | Eosinophils |
| --- | --- | --- | --- | --- | --- |
| FPN2-WS | 0.25X | 0.98 | 0.85 | 0.95 | 0.35 |
| FPN2-WS | 0.5X | **0.98** | 0.86 | 0.97 | 0.45 |
| FPN2-WS | 0.75X | 0.98 | 0.87 | 0.97 | 0.38 |
| FPN2-WS | 1X | 0.98 | 0.87 | **0.97** | 0.46 |
| MRCNN | 0.25X | 0.93 | 0.87 | 0.91 | 0.76 |
| MRCNN | 0.5X | 0.96 | 0.87 | 0.93 | 0.78 |
| MRCNN | 0.75X | 0.97 | **0.9** | 0.93 | 0.77 |
| MRCNN | 1X | 0.97 | 0.9 | 0.94 | **0.8** |

**Table 5).** F1 Scores at IoU[t = 0.7] for the MRCNN and FPN2-WS models trained on augmented ROIs datasets of increasing size. MRCNN and FPN2-WS were trained on augmented ROI image datasets of different sizes from all cell types, as indicated in Fig. 4B. The table indicates the test F1 scores for MRCNN and FPN2-WS object-level segmentation (IoU[t = 0.7]) for each of the 4 cell types in the training conditions examined. The best F1 value for each Cell Type/Model/Training Dataset Size combination is indicated in bold.

| Model | Set Size | MCF10A | HCT116 | U2OS | Eosinophils |
| --- | --- | --- | --- | --- | --- |
| FPN2-WS | 0.125X | 0.97 | 0.81 | 0.88 | 0.41 |
| FPN2-WS | 0.25X | 0.96 | 0.84 | 0.93 | 0.38 |
| FPN2-WS | 0.5X | 0.98 | 0.86 | 0.97 | 0.43 |
| FPN2-WS | 1X | 0.98 | 0.87 | **0.97** | 0.46 |
| FPN2-WS | 2X | **0.99** | 0.86 | 0.97 | 0.44 |
| MRCNN | 0.125X | 0.96 | 0.77 | 0.9 | 0.53 |
| MRCNN | 0.25X | 0.97 | 0.84 | 0.94 | 0.6 |
| MRCNN | 0.5X | 0.97 | 0.88 | 0.94 | 0.78 |
| MRCNN | 1X | 0.97 | 0.9 | 0.94 | **0.8** |
| MRCNN | 2X | 0.98 | **0.91** | 0.96 | 0.68 |

**Table 6).** F1 Scores at IoU[t = 0.7] for the MRCNN and FPN2-WS models trained on differently augmented ROIs datasets. MRCNN and FPN2-WS were trained on ROI image datasets augmented with different strategies, as indicated in Fig. 4C. The table indicates the test F1 scores for MRCNN and FPN2-WS object-level segmentation (IoU[t = 0.7]) for each of the 4 cell types in the training conditions examined. The best F1 value for each Cell Type/Model/Training Dataset Augmentation Strategy combination is indicated in bold.

| Model | Augmentation | MCF10A | HCT116 | U2OS | Eosinophils |
| --- | --- | --- | --- | --- | --- |
| **FPN2-WS** | Full | **0.98** | 0.87 | **0.97** | 0.46 |
| **FPN2-WS** | Min-Blur | 0.98 | 0.86 | 0.97 | 0.37 |
| **FPN2-WS** | Min-Contr | 0.98 | 0.8 | 0.96 | 0.54 |
| **FPN2-WS** | Min-Noise | 0.98 | 0.86 | 0.97 | 0.33 |
| **FPN2-WS** | Min-Scaling | 0.98 | 0.88 | 0.95 | 0.45 |
| **FPN2-WS** | None | 0.98 | 0.83 | 0.96 | 0.53 |
| **MRCNN** | Full | 0.97 | 0.9 | 0.94 | 0.8 |
| **MRCNN** | Min-Blur | 0.97 | 0.88 | 0.95 | 0.75 |
| **MRCNN** | Min-Contr | 0.96 | 0.89 | 0.97 | 0.77 |
| **MRCNN** | Min-Noise | 0.98 | 0.85 | 0.96 | 0.79 |
| **MRCNN** | Min-Scaling | 0.98 | 0.88 | 0.97 | **0.82** |
| **MRCNN** | None | 0.97 | **0.91** | 0.97 | 0.82 |

Having explored a set of model training parameters, we then decided to run replicate training experiments to ensure that the model training regimen was stable and to estimate the variance in inference performance between runs (Fig 4D and Table 7). The results of this set of replicate experiments confirm that training of these models leads to reproducible F1 inference performance, and that initialization with pre-trained weights (i.e., transfer learning) leads to a significant improvement in F1 inference scores at IoU[t = 0.7] with MRCNN and FPN2-WS for several cell types (Fig. 4D and Table 7). In addition, they also indicated that skipping image augmentation before MRCNN training significantly improved F1 scores for HCT116 and primary eosinophils, without leading to a significant drop in performance for MCF10A or U2OS cells (Fig. 4D and Table 7). These results indicate that reducing the number of training cycles and datasets, and skipping image augmentation, are viable options to speed-up the training of these models, without leading to a reduction in their performance.

**Table 7).** F1 Scores at IoU[t = 0.7] for repeated training runs of MRCNN and FPN2-WS models. MRCNN and FPN2-WS were repeatedly trained as indicated in Fig. 4D. The table indicates the mean test F1 scores for MRCNN and FPN2-WS object-level segmentation (IoU[t = 0.7]) for each of the 4 cell types in the training conditions examined. The best mean F1 value for each Cell Type/Model/Training Regimen combination is indicated in bold.

| Model | Initialization | Augmentation | MCF10A | U2OS | HCT116 | Eosinophils |
| --- | --- | --- | --- | --- | --- | --- |
| **FPN2-WS** | Random | None | 0.98 | 0.96 | 0.82 | 0.54 |
| **FPN2-WS** | Random | Full | 0.98 | 0.96 | 0.86 | 0.43 |
| **FPN2-WS** | Pre-initialized | None | 0.98 | 0.96 | 0.84 | 0.53 |
| **FPN2-WS** | Pre-initialized | Full | **0.98** | 0.97 | 0.87 | 0.42 |
| **MRCNN** | Random | None | 0.87 | 0.89 | 0.68 | 0.81 |
| **MRCNN** | Random | Full | 0.24 | 0.12 | 0.21 | 0.42 |
| **MRCNN** | Pre-initialized | None | 0.97 | **0.97** | **0.91** | **0.82** |
| **MRCNN** | Pre-initialized | Full | 0.97 | 0.96 | 0.87 | 0.75 |

### Custom Trained Deep Learning Models Match and Exceed the Nuclear Segmentation Performance of Pre-trained Models

We then benchmarked the nuclear segmentation inference performance of MRCNN and FPN2-WS against Jacobkie, a pre-trained deep learning model for nuclear segmentation (33). Jacobkie has a similar model architecture as FPN2-WS, was trained on a large and diverse dataset of 841 images of nuclei from different sources, was ranked2^nd^ out of about 10,000 submissions in the 2018 Kaggle Data Science Competition, and it has been suggested that this model can be used in inference by end users without pre-training (11). In our experiments using Jacobkie on the test images used in this study, visual comparison of inferred nuclear segmentation masks indicated that, while Jacobkie was generally less precise than MRCNN or FPN2-WS at the pixel segmentation level, it indeed achieved a comparable object segmentation performance for MCF10A and HCT116 cells, and an inferior performance on U2OS or primary eosinophils (Fig. 5A), when compared to the custom trained models. In particular, MRCNN seemed to better generalize to irregular nuclear shapes, such as those of primary eosinophils (Fig. 5A). In line with the qualitative visual results, comparison of F1 segmentation scores for the three segmentation models indicated that MRCNN and FPN2-WS achieved similar or better F1 inference scores than Jacobkie for all cell types at IoU[t = 0.7] (Fig. 5B and Table 8), which is indicative of instance segmentation precision. MRCNN and FPN2-WS showed better F1 scores than Jacobkie at IoU[t = 0.9] which is indicative of segmentation precision at the pixel-level, for MCF10A and U2OS (Fig. 5B and Table 8). As for HCT116 and eosinophils, none of the CNN’s showed F1 scores > 0.5, indicative of a failure of these DL models to perform precise segmentation at the pixel-level of nuclei having a small area relative to the pixel size of the image (Table S1).

**Fig. 5).**
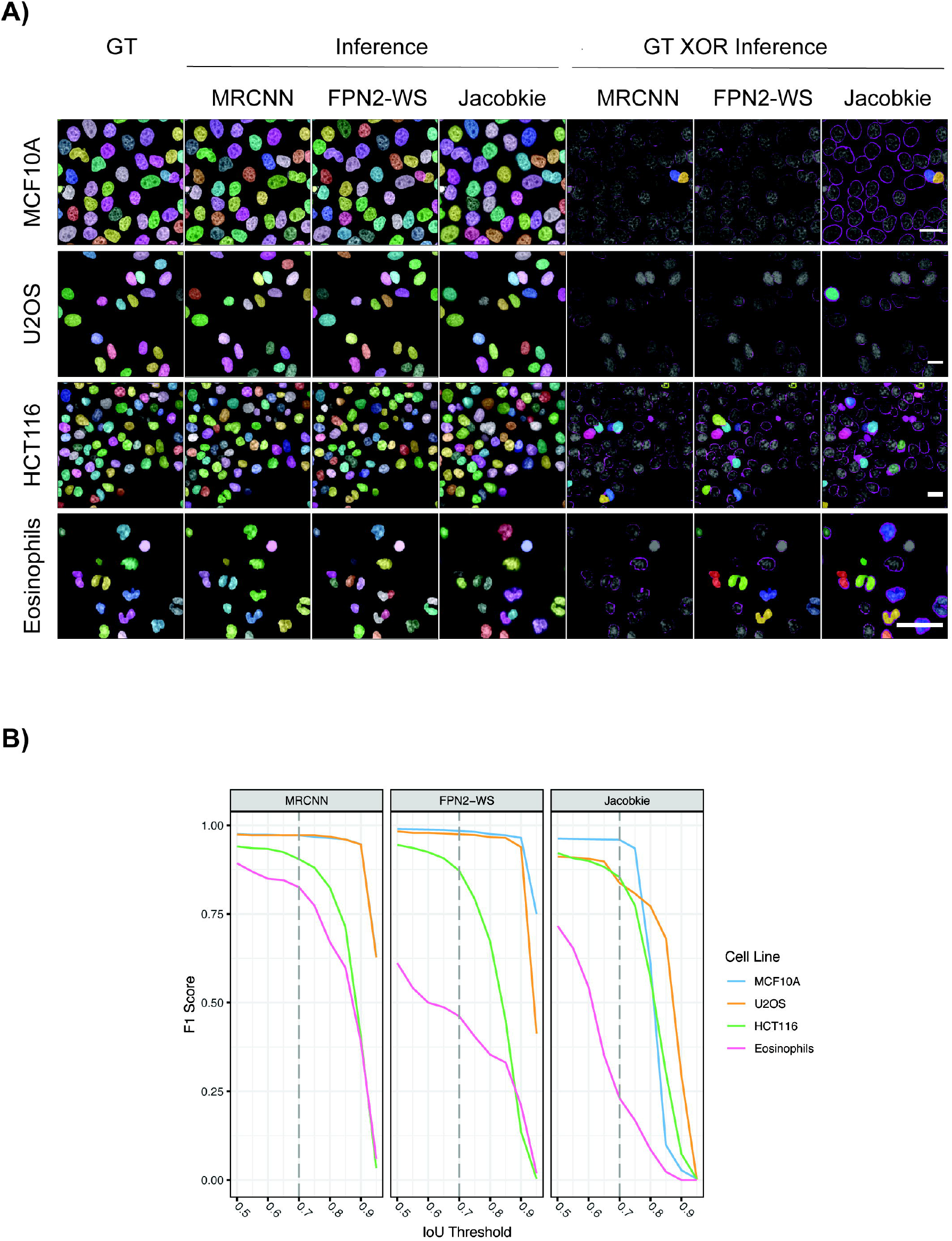
Nuclear segmentation inference performance of the best training strategy for the MRCNN and FPN2-WS model architectures, and comparison to the pre-trained Jacobkie model architecture. **A)** The MRCNN and FPN2-WS models were trained as indicated in Fig. 4D) (None/Pre-initialized for MRCNN, Full Augmentation/Pre-initialized for FPN2-WS, 35,000 bootstrapped ROIs obtained from all the cell types training datasets, 25 cycles). The Jacobkie model was run using the pre-trained weights from the Kaggle Data Science competition (See Materials and Methods for details). The images represent pseudocolored test nuclear labels for the semi-automated ground truth (GT) annotations and for the models test inference results. Additionally, the GT XOR Inference panels represent pseudocolored full labels for false negatives at IoU[t = 0.7]. For true positive and false positive objects at the same threshold, only the difference at the pixel-level between GT and inference labels is shown in magenta. Scale bar: 20 μm. **B)** Line plot of the test F1 Score at increasing IoU thresholds for the MRCNN and FPN2-WS models trained as indicated in A), and for the pre-trained Jacobkie model. IoU[t = 0.7] is indicative of object-level segmentation performance and is indicated in the plot by the dashed grey vertical line.

**Table 8).** F1 Scores at IoU[t = 0.7] and 0.9 for the best training strategy for the MRCNN and FPN2-WS, and comparison with the pre-trained Jacobkie model. MRCNN and FPN2-WS were trained on an image dataset that included all cell types, as indicated in Fig. 5. The table indicates the test F1 scores for MRCNN and FPN2-WS object-level segmentation (IoU[t = 0.7]) and pixel-level segmentation (IoU[t = 0.9]) for each of the 4 cell types. The best F1 value for each IoU/Cell Type combination is indicated in bold.

| Model | MCF10A/0.7 | MCF10A/0.9 | HCT116/0.7 | HCT116/0.9 | U2OS/0.7 | U2OS/0.9 | Eosinophils/0.7 | Eosinophils/0.9 |
| --- | --- | --- | --- | --- | --- | --- | --- | --- |
| FPN2-WS | 0.98 | 0.96 | 0.87 | 0.14 | 0.97 | 0.94 | 0.46 | 0.21 |
| MRCNN | 0.97 | 0.95 | 0.9 | 0.4 | 0.97 | 0.95 | 0.83 | 0.39 |
| Jacobkie | 0.96 | 0.03 | 0.85 | 0.07 | 0.84 | 0.3 | 0.23 | 0 |

### Analysis of the Impact of Semi-annotated Ground Truth Labelling Strategy on Test Segmentation Performance at the Object- and Pixel-level

To rule out the possibility that we introduced a bias in favor of the MRCNN and FPN2-WS in the object-level classification performance by using them as the source of the preliminary GT in the semi-automated labelling strategy, we also tested the segmentation performance of the three CNN models on the test datasets in which the nuclei from the corresponding greyscale images were instead fully manually annotated (Fig S1A, S1B and Table S3). The results of these experiments show that MRCNN and FPN2-WS F1 scores for the comparison of the GT labels generated by the manual vs. the semi-automated method had a minimal difference at IoU[t = 0.7] for all cell types, minimal difference at IoU[t = 0.9] for MCF10A and U2OS, a substantial difference at IoU[t = 0.9] for the HCT116 and eosinophils nuclei (Fig. S1B and Table S3), which can be explained by the fact that the nuclei in these datasets/cell types tend to be very small (Table S1), and thus very likely highly penalized in the F1 score even for small (±1-pixel) errors (See Materials and Methods for details). Overall, these results indicate that semi-automated GT labelling may introduce some bias at the pixel segmentation level that is particularly evident when trying to segment small nuclei, while it has minimal or no effect on the segmentation at the object level. In addition, this also rules out the possibility the Jacobkie model was penalized by the semi-automated labelling procedure, likely because F1 comparison of the manual vs. semi-automated GT labels revealed good concordance for all cell types examined (Fig S2A).

Finally, comparison of the training vs. testing F1 scores for MRCNN and FPN2-WS (Fig. S2B and Table S4) revealed minimal differences between these scores at IoU[t = 0.7] for either model on all 4 cell types. These results suggest that the training regimens described here did not lead to overfitting at the object level. At IoU[t = 0.9], the models did not show overfitting at the pixel level for the MCF10A and U2OS training sets (Fig. S2B and Table S4). For HCT116 and eosinophils at the same threshold, the models performance was too low (i.e., F1 score < 0.5) to make a conclusion about overfitting at the pixel segmentation level, given than the nuclei of these cell types are small relative to the pixel size used during image acquisition (Fig. S2B, Table S1 and S4, and Materials and Methods).

### Dissection of the segmentation errors generated by different nuclear segmentation models

Finally, we dissected the types of errors underlying the nuclear segmentation F1 scores for the three CNN models by examining the frequency of false positives and false negatives (Fig. 5A and S3, and Materials and Methods for details). Additionally, we also examined additional summary statistics, such as over-splitting and merging events (Fig 5A and S3, and Materials and Methods for details), which provide additional information regarding the strengths and weaknesses of each segmentation model (9). The nuclear segmentation error analysis indicated that, while there was no clear pattern in terms of false positive detection among the three models (Fig. S3 A), the Jacobkie model had a tendency to produce a slightly higher rate of false negatives than MRCNN or FPN2-WS (Fig. S3B), possibly because the Jacobkie model was not trained on the image datasets contained in this study. Similarly, FPN2-WS showed a tendency of oversplit single nuclei (Fig. S3C), and MRCNN to merge two or more nuclei (Fig. S3D) when compared to the two other models. Overall, with the exception of the eosinophils, all three models kept the percentage of each type of error below 15%, which is considered acceptable in HCI applications. Moreover, and as manifested in the F1 score curves (Fig 5B), these results suggest that MRCNN might be a better model architecture for the segmentation of heteromorphic, non-convex shaped nuclei when compared to models that incorporate morphological information for segmentation, such as FPN2-WS and Jacobkie (Fig 5A, eosinophils).

Overall, these results confirm that, while in some cases off-the-shelf, pre-trained nuclear segmentation models, such as Jacobkie, can be used for nuclear instance segmentation in inference mode without retraining, optimal nuclear segmentation performance on a subset of cell types with substantially different morphology would ideally uses transfer learning and/or fine-tuning of CNN models on a small number of annotated images.

## Discussion

Machine learning approaches have found wide applications in HCI because they can learn rapidly from the imaging datasets and they generate close to optimal, and parameter-free, solutions to a wide variety of previously non-addressable image processing tasks (6). In particular, in supervised CNN-based machine learning tasks one of the main challenges is to choose appropriate combinations of suitable machine learning model architectures, image pre-processing or post-processing strategies, and model training regimens.

The use of CNN segmentation models pre-trained on large and varied datasets of images has been proposed as a possible solution to these challenges (11)., While very appealing, it is unclear whether this solution would extend to images of nuclei with widely different characteristics such as the ones typically present in large training datasets. A complementary method is to take a CNN-model, possibly pre-trained on a small number of images, and further re-train it for the purpose of fine-tuning using transfer learning on a set of previously unseen nuclei (29). While this strategy would not necessarily guarantee that CNN segmentation models trained in this way will generalize to any previously unseen set of images generated on different instruments or in different laboratories, it might provide a solution similar to what in the machine learning field is known as federated learning, where the CNN models are continuously refined with decentralized data (34). With this framework in place, a lab or an imaging core could either adopt a completely new, untrained CNN model, or start from one of the available domain-specific pre-trained models, and continuously expand their quantitative image segmentation pipelines on images of nuclei from different cell types, if necessary, by iterative cycles of model training. The goal of this study was to test the feasibility of this approach.

We chose a semi-automated ground truth label generation strategy that combines generation of preliminary labels by an algorithm, followed by manual correction of these labels, to reduce the manual annotation burden on the user (Fig 1A). We note that this choice can potentially introduce biases in the CNN models that would be particularly evident in the segmentation of small nuclei or other small cellular objects (such as mitochondria or vesicles). On the one hand, by comparing semi-automated GT labelling with manual GT labelling we observed no difference in F1 scores at IoU[t = 0.9] for any of four cell lines with morphologically vastly distinct cell nuclei (Fig S1, Table S3). On the other, we observed on average a 22% drop in F1 scores for semiautomated labelling GT vs. manual GT labelling at IoU[t = 0.9] for MCF10A and U2OS (Fig S1B, Table S3). As far as small nuclei are concerned (e.g. HCT116 and Eosinophils), while the semi-automated GT labelling methods seemed to introduce a bias that degraded the performance of the trained models by about 70% on average at the pixel level (F1 at IoU[t = 0.9], Fig S1B, Table S3), use of manual GT annotations did not lead to substantially better results in absolute terms (e.g. F1 at IoU[t = 0.9] > 0.5, Fig S1B, Table S3). Overall, these results indicate that semi-automated GT labelling is a viable annotation approach, with the caveat that it can lead to some degradation in precision at the pixel segmentation level. Of course, if the highest level of precision at the pixel level is desired, users should consider manual GT annotation, with the obvious tradeoff of investing a substantially larger amount of hands-on time for this operation. Assisted annotation has already been used both for annotating large biological image datasets for training of CNN-based nuclear segmentation models (11). In addition, similar semiautomated labelling strategies have also been used for very large image datasets of everyday images, such as the Google Open Images Dataset (35).

To test feasibility of training DL models on a limited number of representative images, we used an automated computational pipeline to run a total of 76 supervised model training experiments for CNN tasked to perform segmentation of cell nuclei from fluorescence microscopy images of fixed cells stained with the DNA stain DAPI. To test the usage of nuclear segmentation algorithms in a typical heterogenous laboratory setting, we ran training and inference experiments on a panel of 4 cell types with substantial differences in image acquisition conditions (i.e., magnification and camera binning), nucleus morphology, and degree of confluency. Overall, the FPN2-WS model performed extremely well on 2 of the 4 cell types tested, with or without pre-initialization, and just by training it with images from one cell type (Fig. 2A, Table 2). MRCNN did not perform as well as FPN2-WS in these starting conditions, but it showed comparable or better performance on most cell types when transfer-learning (29) and a training set containing images of the 4 cell types were adopted (Fig. 3A and 3B, Tables 2 and 3). Paradoxically, in the case of MRCNN, adding image augmentation or increasing the number of ROIs used per cell type slightly deteriorated the segmentation performance of the model. While these approaches might sufficient for deep learning-based nuclear segmentation on images acquired in other contexts, our results underscore the importance of testing different training parameters to identify the most important ones, and to potentially reduce the computational load and/or time required for model training. Future efforts to improve the performance of MRCNN on images of nuclei from different cell types, acquired on different fluorescence microscopes, with different microscopy modalities, or different sample types (e.g. histological tissue sections) could take advantage of more advanced transfer learning approaches such as model weight transfusion (30).

Furthermore, comparison of the performance of MRCNN and FPN2-WS trained on our dataset with Jacobkie, a state-of-the-art pre-trained CNN model (11), indicates that using pre-trained models is an effective, lower-effort trade-off on images of nuclei of certain cell types, but that training and testing of CNN models on the images of interest becomes necessary if the final goal is to obtain higher pixel-level precision, such as in the case of MCF10A and U2OS cells (Fig 5A and 5B, Table 8), or if the nuclei in question are substantially different from the ones present in the original training set, such as primary eosinophils (Fig 5A and 5B, Table 8). As a caveat, we also note that if the size of the nuclei is small relative to the pixel size of the images, such as in the case of HCT116 and eosinophils (Table S1), neither training DL models *ex-novo,* nor using a state-of-the art out of the box DL model, seems to achieve a satisfactory performance at the pixel level (Fig 5A and 5B, Table 8). Other potentially useful applications for the use of transfer learning in nuclear segmentation might include the use of other fluorescence markers to identify nuclei in live cell experiments, such as the core histone H2B-GFP or the DNA stain SiR-DNA, and to more precisely segment nuclei in cell types that tend to grow in patches/colonies, such as iPS and mESC cells, where nuclei are packed tightly together. Moreover, when we compared the out-of-the box inference runtime of MRCNN versus Jacobkie, the MRCNN model was 14-fold faster than Jacobkie (data not shown). Of course, in the end, users will need to carefully evaluate the tradeoffs involved in using out-of-the box models, which provide ease of use and but potentially lower accuracy on hard to segment or nuclei with an unusual morphology, versus adopting the transfer learning strategy proposed, which involve substantial computational work for model training to achieve higher segmentation performance in these cases. Ultimately, we see these two approaches not as mutually exclusive, but rather as complementary. To this end, we recommend that end-users first test the inference performance of a pre-trained state-of-the-art nuclear segmentation model such as Jacobkie on the images of interest, and, if these results are not satisfactory, then edit the preliminary nuclear labels and fine-tune the pre-trained model by transfer learning to improve its performance.

We hope that the training pipeline described here, and this practical set of guidelines, will help researchers to test novel model architectures and novel image pre- and postprocessing steps aimed at the segmentation of nuclei and other biological structures from microscopy images. In the future, we expect it will be feasible to expand this set of observations to 3D images of nuclei, and to the segmentation of other cellular organelles and sub-structures, such as mitochondria, vesicles, or nuclear bodies.

## Acknowledgements

The authors thank the CBIIT Server Team, NCI, National Institutes of Health (NIH). This work utilized the computational resources of the NIH HPC Biowulf cluster (http://hpc.nih.gov). Also, we would like to thank Murat Bilgel for contributing to the MRCNN postprocessing implementation, and Andrew Weisman for contributing to the image augmentation wrapper. We thank Amy Klion and Michelle Makiya at the National Institute of Allergy and Infectious Diseases for providing primary human eosinophils. This research was supported by funding from the Intramural Research Program of the National Cancer Institute, Center for Cancer Research Project Num. 1ZICBC011567-01, the National Institute of Allergy and Infectious Diseases, and the National Institute of Arthritis and Musculoskeletal and Skin Diseases, all at the National Institutes of Health (NIH). All the authors declare no conflict of interest.

## Author Contributions

GZ, PRG and GP conceptualized and designed the project; GZ and JK refactored the code for the MRCNN model; PRG wrote the code for the FPN2-WS model; GZ, PRG, and KL wrote and executed the nuclear segmentation training pipeline; LO, SS, MG, JS, IF, LMF, TM, and GP provided the training images dataset; GZ and GP analyzed the results of the experiments; GZ and GP wrote the manuscript; all authors read and revised the manuscript.

## Supplementary Figures and Tables Legend

**Fig. S1) Comparison of the inference performance at testing for the MRCNN and FPN2-WS, and for the Jacobkie model using testing labels that were either semiautomated or manually annotated. A)** MRCNN and FPN2-WS were trained as indicated in Fig. 5), and Jacobkie model was run using the pre-trained weights from the Kaggle Data Science competition (See Materials and Methods for details). The images represent pseudocolored test nuclear labels for the manual ground truth (GT) annotations and for the model inference results, as shown in Fig 5A. Additionally, the GT XOR Inference panels represent pseudocolored full labels for false negatives at an Intersection over Union (IoU) threshold of 0.7. For true positive and false positive objects at the same threshold, only the difference at the pixel-level between the manual GT and inference labels is shown in magenta. Scale bar: 20 μm. **B)** Line plots of the test F1 Score values obtained using either the manual GT labels or the semi-automated GT labels for the same greyscale images, respectively, at increasing IoU thresholds for the MRCNN and FPN2-WS models trained as indicated in A), and for the pre-trained Jacobkie model. IoU[t = 0.7] is indicated by the dashed grey vertical line.

**Fig. S2) Quality control measurements for the semi-automated GT label annotation strategy and for model overfitting. A)** Line plots of the F1 Score values at increasing IoU thresholds comparing the manual GT labels with the semi-automated GT labels for the same greyscale images. IoU[t = 0.7] is indicated by the dashed grey vertical line in the plot. **B)** Line plots of the training F1 Score vs. the testing F1 Score at increasing IoU thresholds using the semi-automated GT labels for the training and testing datasets of greyscale images, for the MRCNN and FPN2-WS models trained as indicated in Fig 5. IoU[t = 0.7] is indicated by the dashed grey vertical line.

**Fig. S3) Analysis of the segmentation errors types generated by the MRCNN, FPN2-WS, and Jacobkie models.** Bar plots of the percentage of errors types generated by the nuclear segmentation models at test inference (As shown in Fig. 5). **A)** False positives. **B)** False Negatives. **C)** Merges. **D)** Oversplitting. Notice that each the y-axis of each plot facet has been scaled for each error type and each cell line tested, to make possible an internal comparison of the three models on the same cell line. Also, notice that, by definition, certain types of errors can be counted two or more times (See Materials and Methods for details). Thus, the sum of error percentages for each model and cell line does not necessarily sum to 100%.

**Table S1) Characteristics of the training and testing image datasets used in this work.**

**Table S2) List of all model training conditions with references to the other Figures and Supplementary Figures presented in this work.**

**Table S3) F1 Scores at IoU[t = 0.7] and IoU[t = 0.9] for the inference performance comparison between semi-automated and manually annotated testing labels.** MRCNN and FPN2-WS were trained on an image dataset that included all cell types, as indicated in Fig. 5 and S1. The table indicates the test F1 scores for MRCNN and FPN2-WS object-level segmentation (IoU[t = 0.7]) and pixel-level segmentation (IoU[t = 0.9]) for each of the 4 cell types. The best test F1 value for each IoU/Cell Type combination is indicated in bold.

**Table S4) Training and Testing F1 Scores at IoU[t = 0.7] and IoU[t = 0.9].** MRCNN and FPN2-WS were trained on an image dataset that included all cell types, as indicated in Fig. 5. The table indicates the training and testing F1 scores for MRCNN and FPN2-WS object-level segmentation (IoU[t = 0.7]) and pixel-level segmentation (IoU[t = 0.9]) for each of the 4 cell types. The best training or testing F1 value for each IoU/Cell Type combination is indicated in bold.

