## Supplementary figures and images for "A Deep Learning Pipeline for Nucleus Segmentation"

### Fig. S1

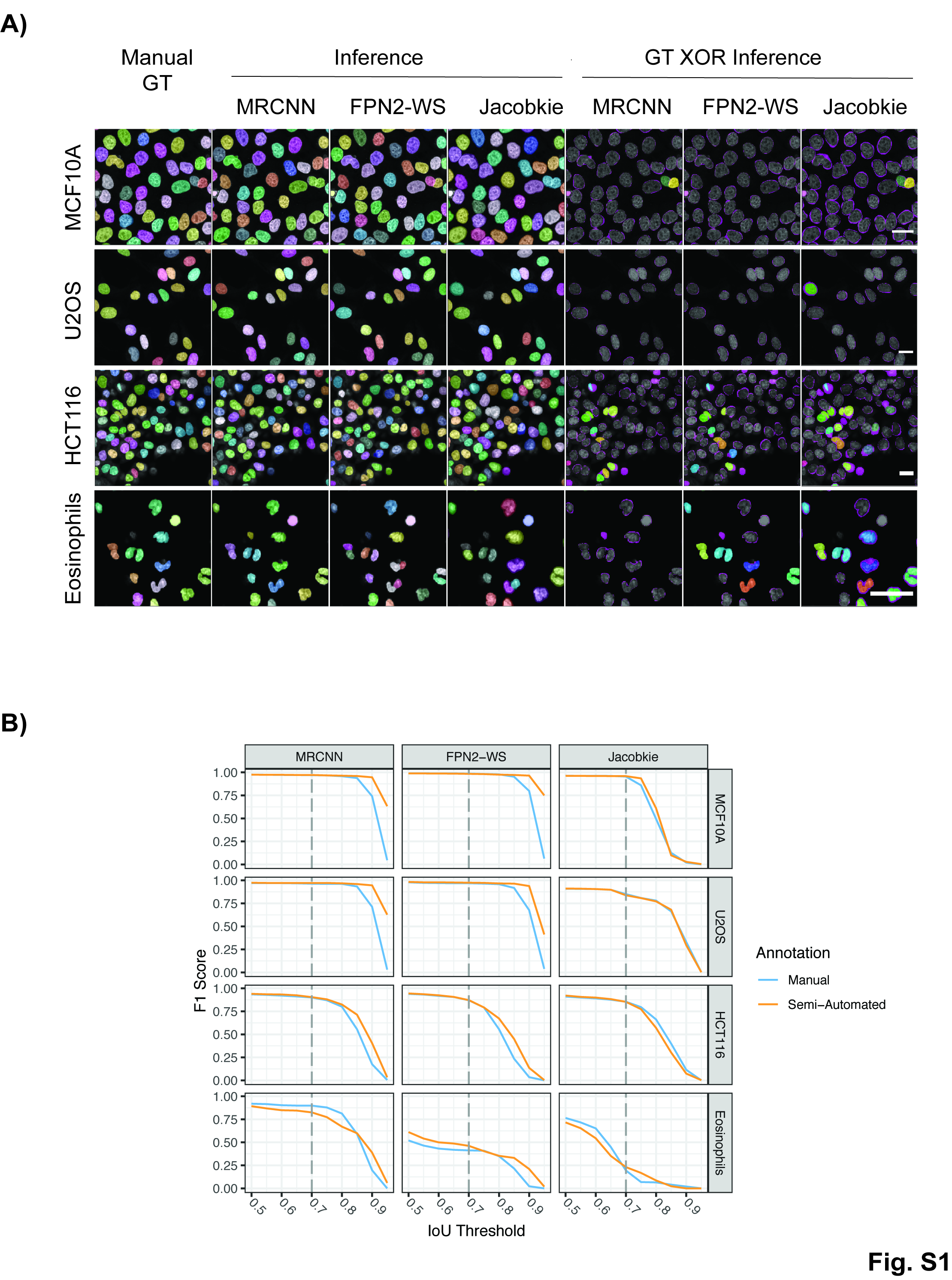

### Fig. S2

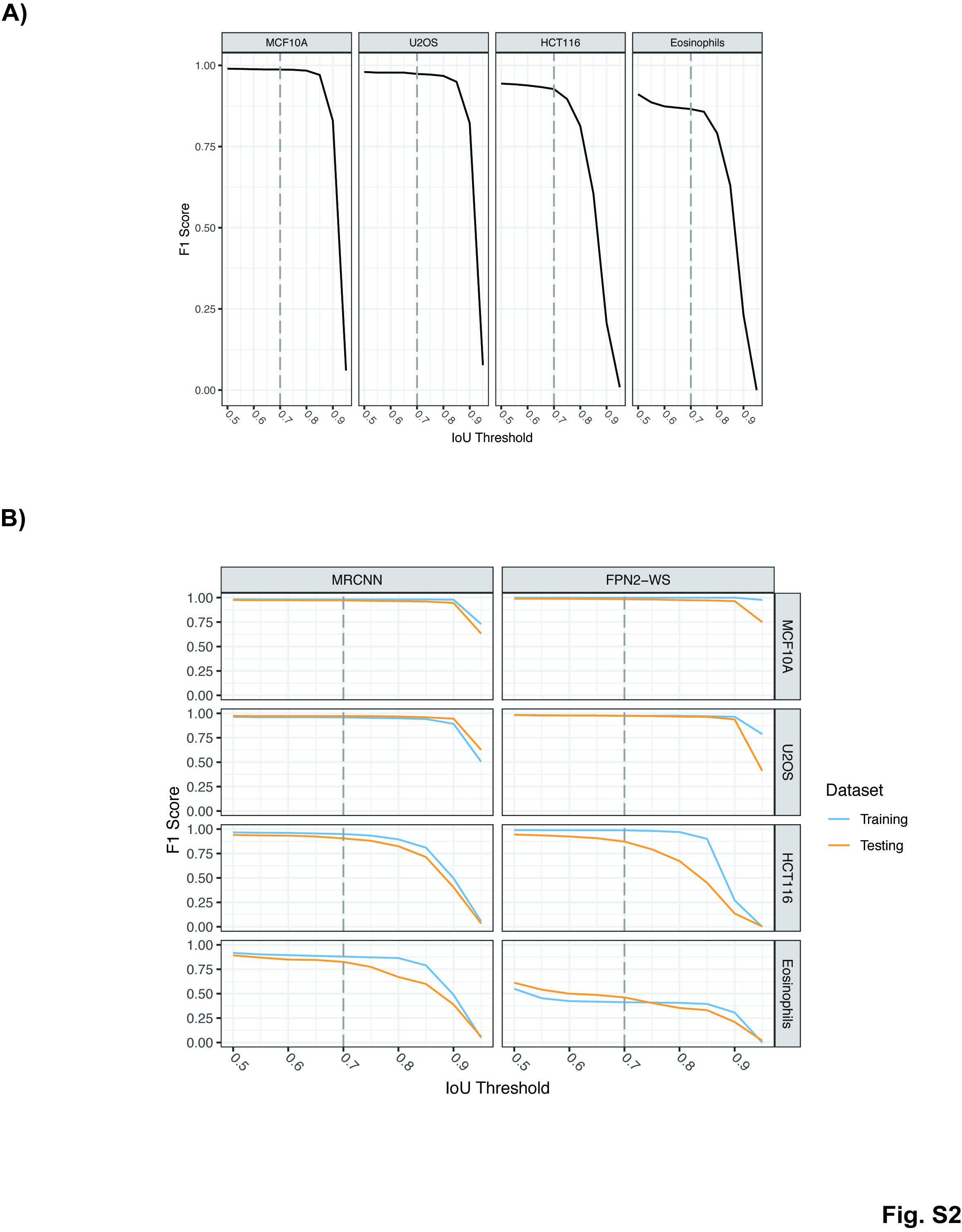

### Fig. S3

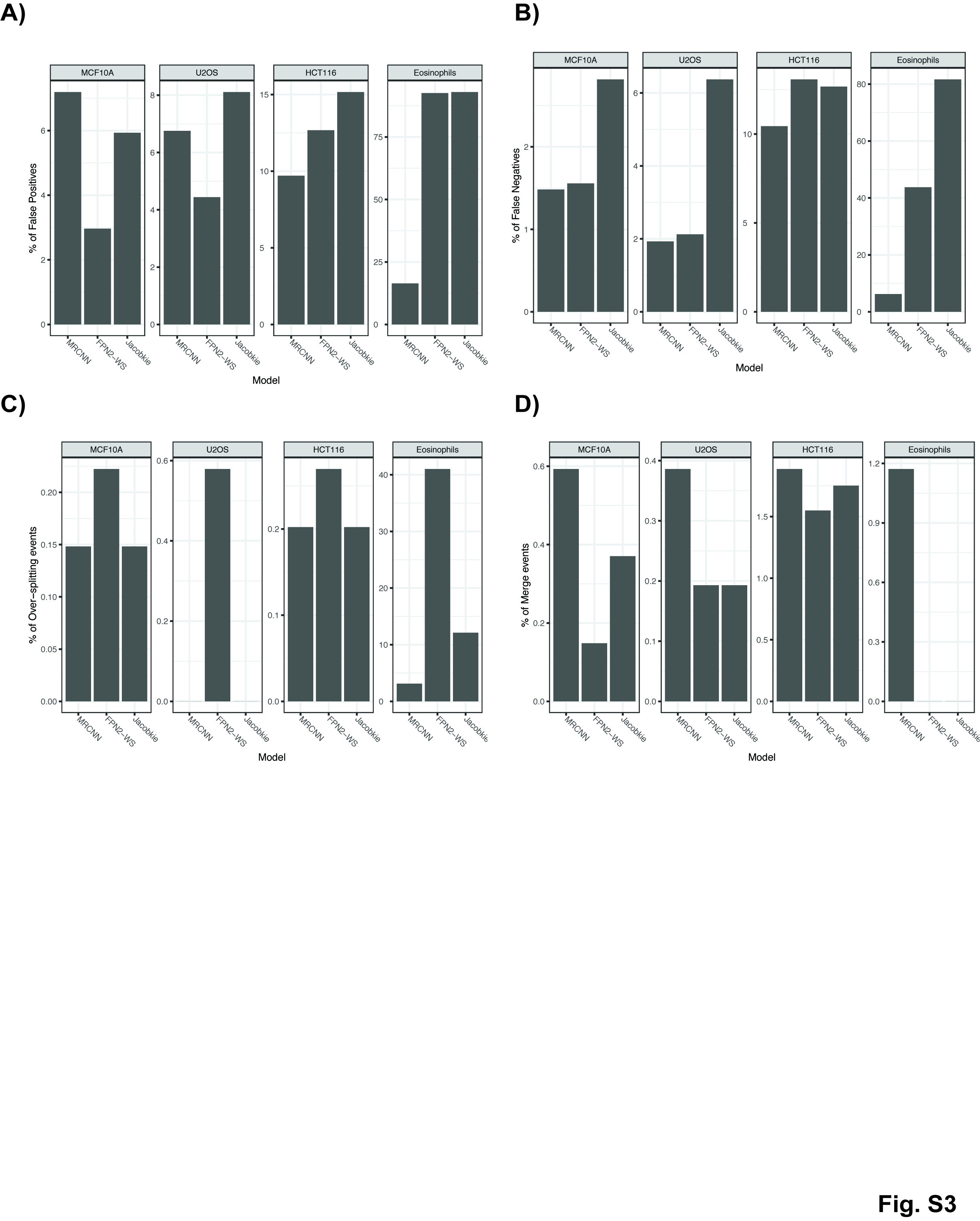
