## Supplementary material for "A Deep Learning Pipeline for Nucleus Segmentation": Table S1

| **Dataset Name** | **Number FOVs** | **FOV Size - px** | **Objective** | **Binning** | **XY Px Size - micron** | **Number of Nuclei** | **Area Mean - px** | **Area SD - px** | **Dataset Type** | **GT Labelling** |
| --- | --- | --- | --- | --- | --- | --- | --- | --- | --- | --- |
| **MCF10A_Original** | **7** | **1078 x 1278** | **60X** | **2 x 2** | **0.216** | **1225** | **2606.10** | **855.10** | **Train** | **Semi-Auto** |
| **MCF10A_Biological** | 7 | 1078 x 1278 | 60X | 2 x 2 | 0.216 | 1349 | 2605.57 | 738.13 | Test | Manual |
| **MCF10A_Biological** | 7 | 1078 x 1278 | 60X | 2 x 2 | 0.216 | 1382 | 2575.46 | 828.18 | Test | Semi-Auto |
| **HCT-116_Original** | 1 | 1080 x 1280 | 20X | 2 x 2 | 0.650 | 1710 | 299.05 | 91.43 | Train | Semi-Auto |
| **HCT-116_Biological** | 1 | 1080 x 1280 | 20X | 2 x 2 | 0.650 | 1484 | 300.31 | 97.69 | Test | Manual |
| **HCT-116_Biological** | 1 | 1080 x 1280 | 20X | 2 x 2 | 0.650 | 1488 | 299.10 | 96.38 | Test | Semi-Auto |
| **U2OS_Original** | 1 | 2158 x 2558 | 20X | 1 x 1 | 0.325 | 782 | 2175.55 | 752.44 | Train | Semi-Auto |
| **U2OS_Technical** | 1 | 2158 x 2558 | 20X | 1 x 1 | 0.325 | 518 | 1991.52 | 663.17 | Test | Manual |
| **U2OS_Technical** | 1 | 2158 x 2558 | 20X | 1 x 1 | 0.325 | 529 | 1926.78 | 687.21 | Test | Semi-Auto |
| **Eosinophils_Original** | 1 | 1080 x 1280 | 60X | 2 x 2 | 0.216 | 295 | 562.38 | 183.31 | Train | Semi-Auto |
| **Eosinophils_Technical** | 1 | 1080 x 1280 | 60X | 2 x 2 | 0.216 | 256 | 566.41 | 157.88 | Test | Manual |
| **Eosinophils_Technical** | 1 | 1080 x 1280 | 60X | 2 x 2 | 0.216 | 270 | 528.01 | 185.87 | Test | Semi-Auto |
