## Supplementary material for "A Deep Learning Pipeline for Nucleus Segmentation": Table S2

| **Figure** | **Experiment** | **Training Dataset(s)** | **Number of ROIS for Training** | **Augmentation** | **Number of Cycles** | **Model** | **Initialization Weights** |
| --- | --- | --- | --- | --- | --- | --- | --- |
| **2B** | 3 | MCF10A | 35,000 | Full | 25 | MRCNN | Random |
|  | 4 | MCF10A | 35,000 | Full | 25 | FPN2-WS | Random |
| **3A** | 1 | MCF10A | 35,000 | Full | 25 | MRCNN | COCO |
|  | 3 | MCF10A | 35,000 | Full | 25 | MRCNN | Random |
|  | 4 | MCF10A | 35,000 | Full | 25 | FPN2-WS | Random |
|  | 5 | MCF10A | 35,000 | Full | 25 | FPN2-WS | ImageNet |
| **3B** | 1 | MCF10A | 35,000 | Full | 25 | MRCNN | COCO |
|  | 5 | MCF10A | 35,000 | Full | 25 | FPN2-WS | ImageNet |
|  | 6 | MCF10A,HCT116,U2OS,Eosinophils | 35,000 | Full | 25 | MRCNN | COCO |
|  | 10 | MCF10A,HCT116,U2OS,Eosinophils | 35,000 | Full | 25 | FPN2-WS | ImageNet |
|  | 29 | MCF10A_HCT116 | 35,000 | Full | 25 | MRCNN | COCO |
|  | 30 | MCF10A_HCT116_U2OS | 35,000 | Full | 25 | MRCNN | COCO |
|  | 65 | MCF10A_HCT116 | 35,000 | Full | 25 | FPN2-WS | ImageNet |
|  | 66 | MCF10A_HCT116_U2OS | 35,000 | Full | 25 | FPN2-WS | ImageNet |
| **4A** | 6 | MCF10A,HCT116,U2OS,Eosinophils | 35,000 | Full | 25 | MRCNN | COCO |
|  | 10 | MCF10A,HCT116,U2OS,Eosinophils | 35,000 | Full | 25 | FPN2-WS | ImageNet |
|  | 12 | MCF10A,HCT116,U2OS,Eosinophils | 35,000 | Full | 14 | FPN2-WS | ImageNet |
|  | 16 | MCF10A,HCT116,U2OS,Eosinophils | 35,000 | Full | 6.25 | MRCNN | COCO |
|  | 17 | MCF10A,HCT116,U2OS,Eosinophils | 35,000 | Full | 12.5 | MRCNN | COCO |
|  | 18 | MCF10A,HCT116,U2OS,Eosinophils | 35,000 | Full | 18.75 | MRCNN | COCO |
|  | 67 | MCF10A,HCT116,U2OS,Eosinophils | 35,000 | Full | 20 | FPN2-WS | ImageNet |
|  | 68 | MCF10A,HCT116,U2OS,Eosinophils | 35,000 | Full | 8 | FPN2-WS | ImageNet |
| **4B** | 6 | MCF10A,HCT116,U2OS,Eosinophils | 35,000 | Full | 25 | MRCNN | COCO |
|  | 10 | MCF10A,HCT116,U2OS,Eosinophils | 35,000 | Full | 25 | FPN2-WS | ImageNet |
|  | 11 | MCF10A,HCT116,U2OS,Eosinophils | 70,000 | Full | 25 | FPN2-WS | ImageNet |
|  | 19 | MCF10A,HCT116,U2OS,Eosinophils | 17,500 | Full | 25 | MRCNN | COCO |
|  | 20 | MCF10A,HCT116,U2OS,Eosinophils | 70,000 | Full | 25 | MRCNN | COCO |
|  | 45 | MCF10A,HCT116,U2OS,Eosinophils | 8,750 | Full | 25 | MRCNN | COCO |
|  | 46 | MCF10A,HCT116,U2OS,Eosinophils | 4,375 | Full | 25 | MRCNN | COCO |
|  | 47 | MCF10A,HCT116,U2OS,Eosinophils | 17,500 | Full | 25 | FPN2-WS | ImageNet |
|  | 48 | MCF10A,HCT116,U2OS,Eosinophils | 4,375 | Full | 25 | FPN2-WS | ImageNet |
|  | 49 | MCF10A,HCT116,U2OS,Eosinophils | 17,500 | Full | 25 | FPN2-WS | ImageNet |
| **4C** | 6 | MCF10A,HCT116,U2OS,Eosinophils | 35,000 | Full | 25 | MRCNN | COCO |
|  | 10 | MCF10A,HCT116,U2OS,Eosinophils | 35,000 | Full | 25 | FPN2-WS | ImageNet |
|  | 22 | MCF10A,HCT116,U2OS,Eosinophils | 35,000 | Min_Contr | 25 | MRCNN | COCO |
|  | 23 | MCF10A,HCT116,U2OS,Eosinophils | 35,000 | Min_Blur | 25 | MRCNN | COCO |
|  | 24 | MCF10A,HCT116,U2OS,Eosinophils | 35,000 | Min_Noise | 25 | MRCNN | COCO |
|  | 25 | MCF10A,HCT116,U2OS,Eosinophils | 35,000 | Min_Scaling | 25 | MRCNN | COCO |
|  | 27 | MCF10A,HCT116,U2OS,Eosinophils | 35,000 | None | 25 | MRCNN | COCO |
|  | 62 | MCF10A,HCT116,U2OS,Eosinophils | 35,000 | None | 25 | FPN2-WS | ImageNet |
|  | 69 | MCF10A,HCT116,U2OS,Eosinophils | 35,000 | Min_Contr | 25 | FPN2-WS | ImageNet |
|  | 70 | MCF10A,HCT116,U2OS,Eosinophils | 35,000 | Min_Blur | 25 | FPN2-WS | ImageNet |
|  | 71 | MCF10A,HCT116,U2OS,Eosinophils | 35,000 | Min_Noise | 25 | FPN2-WS | ImageNet |
|  | 72 | MCF10A,HCT116,U2OS,Eosinophils | 35,000 | Min_Scaling | 25 | FPN2-WS | ImageNet |
| **4D** | 6 | MCF10A,HCT116,U2OS,Eosinophils | 35,000 | Full | 25 | MRCNN | COCO |
|  | 8 | MCF10A,HCT116,U2OS,Eosinophils | 35,000 | Full | 25 | MRCNN | Random |
|  | 10 | MCF10A,HCT116,U2OS,Eosinophils | 35,000 | Full | 25 | FPN2-WS | ImageNet |
|  | 26 | MCF10A,HCT116,U2OS,Eosinophils | 35,000 | Full | 25 | MRCNN | COCO |
|  | 32 | MCF10A,HCT116,U2OS,Eosinophils | 35,000 | Full | 25 | MRCNN | COCO |
|  | 33 | MCF10A,HCT116,U2OS,Eosinophils | 35,000 | Full | 25 | MRCNN | COCO |
|  | 34 | MCF10A,HCT116,U2OS,Eosinophils | 35,000 | Full | 25 | MRCNN | Random |
|  | 35 | MCF10A,HCT116,U2OS,Eosinophils | 35,000 | Full | 25 | MRCNN | Random |
|  | 36 | MCF10A,HCT116,U2OS,Eosinophils | 35,000 | Full | 25 | MRCNN | Random |
|  | 37 | MCF10A,HCT116,U2OS,Eosinophils | 35,000 | None | 25 | MRCNN | COCO |
|  | 38 | MCF10A,HCT116,U2OS,Eosinophils | 35,000 | None | 25 | MRCNN | COCO |
|  | 39 | MCF10A,HCT116,U2OS,Eosinophils | 35,000 | None | 25 | MRCNN | COCO |
|  | 40 | MCF10A,HCT116,U2OS,Eosinophils | 35,000 | None | 25 | MRCNN | Random |
|  | 41 | MCF10A,HCT116,U2OS,Eosinophils | 35,000 | None | 25 | MRCNN | Random |
|  | 42 | MCF10A,HCT116,U2OS,Eosinophils | 35,000 | None | 25 | MRCNN | Random |
|  | 43 | MCF10A,HCT116,U2OS,Eosinophils | 35,000 | None | 25 | MRCNN | Random |
|  | 50 | MCF10A,HCT116,U2OS,Eosinophils | 35,000 | Full | 25 | FPN2-WS | Random |
|  | 51 | MCF10A,HCT116,U2OS,Eosinophils | 35,000 | Full | 25 | FPN2-WS | Random |
|  | 52 | MCF10A,HCT116,U2OS,Eosinophils | 35,000 | Full | 25 | FPN2-WS | Random |
|  | 53 | MCF10A,HCT116,U2OS,Eosinophils | 35,000 | Full | 25 | FPN2-WS | Random |
|  | 54 | MCF10A,HCT116,U2OS,Eosinophils | 35,000 | None | 25 | FPN2-WS | Random |
|  | 55 | MCF10A,HCT116,U2OS,Eosinophils | 35,000 | None | 25 | FPN2-WS | Random |
|  | 56 | MCF10A,HCT116,U2OS,Eosinophils | 35,000 | None | 25 | FPN2-WS | Random |
|  | 57 | MCF10A,HCT116,U2OS,Eosinophils | 35,000 | None | 25 | FPN2-WS | Random |
|  | 58 | MCF10A,HCT116,U2OS,Eosinophils | 35,000 | Full | 25 | FPN2-WS | ImageNet |
|  | 59 | MCF10A,HCT116,U2OS,Eosinophils | 35,000 | Full | 25 | FPN2-WS | ImageNet |
|  | 60 | MCF10A,HCT116,U2OS,Eosinophils | 35,000 | Full | 25 | FPN2-WS | ImageNet |
|  | 61 | MCF10A,HCT116,U2OS,Eosinophils | 35,000 | None | 25 | FPN2-WS | ImageNet |
|  | 62 | MCF10A,HCT116,U2OS,Eosinophils | 35,000 | None | 25 | FPN2-WS | ImageNet |
|  | 63 | MCF10A,HCT116,U2OS,Eosinophils | 35,000 | None | 25 | FPN2-WS | ImageNet |
|  | 64 | MCF10A,HCT116,U2OS,Eosinophils | 35,000 | None | 25 | FPN2-WS | ImageNet |
|  | 73 | MCF10A,HCT116,U2OS,Eosinophils | 35,000 | Full | 25 | MRCNN | Random |
|  | 74 | MCF10A,HCT116,U2OS,Eosinophils | 35,000 | Full | 25 | MRCNN | Random |
|  | 75 | MCF10A,HCT116,U2OS,Eosinophils | 35,000 | Full | 25 | MRCNN | Random |
|  | 76 | MCF10A,HCT116,U2OS,Eosinophils | 35,000 | Full | 25 | MRCNN | Random |
|  | 77 | MCF10A,HCT116,U2OS,Eosinophils | 35,000 | Full | 25 | MRCNN | Random |
|  | 78 | MCF10A,HCT116,U2OS,Eosinophils | 35,000 | Full | 25 | MRCNN | Random |
|  | 79 | MCF10A,HCT116,U2OS,Eosinophils | 35,000 | Full | 25 | MRCNN | Random |
|  | 80 | MCF10A,HCT116,U2OS,Eosinophils | 35,000 | Full | 25 | MRCNN | Random |
|  | 81 | MCF10A,HCT116,U2OS,Eosinophils | 35,000 | Full | 25 | MRCNN | Random |
|  | 82 | MCF10A,HCT116,U2OS,Eosinophils | 35,000 | Full | 25 | MRCNN | Random |
|  | 83 | MCF10A,HCT116,U2OS,Eosinophils | 35,000 | Full | 25 | MRCNN | Random |
| **5B** | 10 | MCF10A,HCT116,U2OS,Eosinophils | 35,000 | Full | 25 | FPN2-WS | ImageNet |
|  | 37 | MCF10A,HCT116,U2OS,Eosinophils | 35,000 | None | 25 | MRCNN | COCO |
|  | 44 | BBBC038 | NA | NA | NA | Jacobkie | Jacobkie |
| **S1B** | 10 | MCF10A,HCT116,U2OS,Eosinophils | 35,000 | Full | 25 | FPN2-WS | ImageNet |
|  | 37 | MCF10A,HCT116,U2OS,Eosinophils | 35,000 | None | 25 | MRCNN | COCO |
|  | 44 | BBBC038 | NA | NA | NA | Jacobkie | Jacobkie |
| **S2B** | 10 | MCF10A,HCT116,U2OS,Eosinophils | 35,000 | Full | 25 | FPN2-WS | ImageNet |
|  | 37 | MCF10A,HCT116,U2OS,Eosinophils | 35,000 | None | 25 | MRCNN | COCO |

**Table S2**
