## Supplementary material for "A Deep Learning Pipeline for Nucleus Segmentation": Table S3

| **Model** | **GT Annotation** | **MCF10A/0.7** | **MCF10A/0.9** | **HCT116/0.7** | **HCT116/0.9** | **U2OS/0.7** | **U2OS/0.9** | **Eosinophils/0.7** | **Eosinophils/0.9** |
| --- | --- | --- | --- | --- | --- | --- | --- | --- | --- |
| **FPN2-WS** | Manual | **0.99** | 0.8 | 0.87 | 0.03 | 0.97 | 0.68 | 0.41 | 0.02 |
| **FPN2-WS** | SemiAutomated | 0.98 | **0.96** | 0.87 | 0.14 | **0.97** | 0.94 | 0.46 | 0.21 |
| **MRCNN** | Manual | 0.97 | 0.74 | 0.9 | 0.18 | 0.96 | 0.71 | **0.9** | 0.2 |
| **MRCNN** | SemiAutomated | 0.97 | 0.95 | **0.9** | **0.4** | 0.97 | **0.95** | 0.83 | **0.39** |
| **Jacobkie** | Manual | 0.95 | 0.02 | 0.85 | 0.12 | 0.85 | 0.33 | 0.19 | 0.02 |
| **Jacobkie** | SemiAutomated | 0.96 | 0.03 | 0.85 | 0.07 | 0.84 | 0.3 | 0.23 | 0 |
