## Supplementary material for "A Deep Learning Pipeline for Nucleus Segmentation": Table S4

| **Model** | **Inference Set** | **MCF10A/0.7** | **MCF10A/0.9** | **HCT116/0.7** | **HCT116/0.9** | **U2OS/0.7** | **U2OS/0.9** | **Eosinophils/0.7** | **Eosinophils/0.9** |
| --- | --- | --- | --- | --- | --- | --- | --- | --- | --- |
| **FPN2-WS** | Training | **1** | **1** | **0.99** | 0.27 | **0.98** | **0.97** | 0.41 | 0.31 |
| **FPN2-WS** | Testing | 0.98 | 0.96 | 0.87 | 0.14 | 0.97 | 0.94 | 0.46 | 0.21 |
| **MRCNN** | Training | 0.98 | 0.98 | 0.95 | **0.5** | 0.96 | 0.89 | **0.88** | **0.49** |
| **MRCNN** | Testing | 0.97 | 0.95 | 0.9 | 0.4 | 0.97 | 0.95 | 0.83 | 0.39 |

**Table S4**
